# Mis-regulation of the Nucleoporins 98 and 96 lead to defects in protein synthesis that promote hallmarks of tumorigenesis

**DOI:** 10.1101/2021.08.02.454839

**Authors:** Ajai J. Pulianmackal, Kiriaki Kanakousaki, Kerry A. Flegel, Olga G. Grushko, Ella Gourley, Emily Rozich, Laura A. Buttitta

**Author notes:** These authors contributed equally (Ajai J. Pulianmackal, Kiriaki Kanakousaki).

## Abstract

The Nucleoporin 98KD (Nup98) is one of the most promiscuous translocation partners in hematological malignancies, contributing to at least 31 different truncation-fusion proteins. To date, nearly all disease models of Nup98 translocations involve ectopic expression of transgenes recapitulating the fusion protein under study, leaving the endogenous *Nup98* loci unperturbed. Overlooked in these approaches is that translocation leads to the loss of one copy of normal Nup98 in addition to the loss of Nup96 – a second Nucleoporin encoded within the same mRNA and reading frame as Nup98. Nup98 and 96 are also mutated in a number of other cancer types and are located near a tumor suppressor region known to be epigenetically silenced, suggesting that their disruption is not limited to blood cancers. We found that reducing Nup98-96 function via an RNAi approach in *Drosophila melanogaster* (where the Nup98-96 shared mRNA and reading frame gene structure is conserved) de-regulates the cell cycle. We find evidence of over-proliferation in Nup98-96 deficient tissues, counteracted by elevated apoptosis and aberrant Wingless and JNK signaling associated with chronic wound healing. When the knockdown of Nup98-96 is combined with inhibition of apoptosis, we see synergism leading to dramatic tissue overgrowth, consistent with a tumor-suppressor function for endogenous Nup98 and 96. To understand how growth and proliferation become mis-regulated when Nup98-96 levels are reduced, we performed RNAseq and uncovered a gene expression signature consistent with defects in ribosome biogenesis. We found that reducing Nup 98 and 96 function limits nuclear export of the ribosome component RpL10A, leading to defects in protein synthesis. Defects in protein synthesis are sufficient to trigger JNK signaling that contributes to compensatory proliferation and hallmarks of tumorigenesis when apoptosis is inhibited. Based upon our data, we suggest that the partial loss of Nup98 and Nup96 function in translocations could de-regulate protein synthesis leading to stress signaling that cooperates with other mutations in cancer to promote tumorigenesis.

**Highlights:**

- Compromising Nups 98 and 96 triggers cell death and compensatory proliferation via JNK signaling that becomes tumorigenic when apoptosis is blocked
- Reducing Nup 98 and 96 function limits nuclear export of the ribosome stalk component RpL10A, leading to defects in protein synthesis which cause stress signaling via JNK.
- Reduced protein synthesis coupled with increased JNK signaling, paradoxically leads to more rapid proliferation with a gene expression signature that resembles a chronic wounding response.
- Overexpression of Nup98, which occurs in oncogenic fusions, leads to similar defects in protein synthesis and JNK activation.

## Introduction

Communication between the nucleus and cytoplasm occurs through nuclear pore complexes (NPCs), which are composed of highly conserved proteins termed Nucleoporins (Nups). Mutations in several Nups are associated with cancer, including loss-of-function mutations and translocations (Simon and Rout, 2014). Of the Nups associated with translocations, Nup98 is the most promiscuous (Lam and Aplan, 2001; Simon and Rout, 2014).

Nup98 function has been difficult to examine because the gene locus for Nup98 encodes for two essential Nucleoporins, Nup98 and Nup96, which derive from an autocatalytic cleavage of a larger Nup98-96 polypeptide with Nup98 located at the amino terminus (Fontoura et al., 1999; Rosenblum and Blobel, 1999). However, a shorter Nup98 only transcript is also produced by the locus via alternative splicing (Fontoura et al., 1999). Nup98 is a peripheral Nup, found both in nuclear pores and in the nucleoplasm (Griffis et al., 2002). It contains FG (Phenylalanine-Glycine) and GLFG repeats in its N-terminal region that allow Nup98 to interact with different nuclear transport receptors (Bachi et al., 2000; Moroianu et al., 1995) during nucleocytoplasmic shuttling, and it has a role in regulating gene transcription (Capelson et al., 2010; Kalverda et al., 2010). In contrast, Nup96 is a core scaffold protein; it is stably localized at NPC and is part of the core Nup107-160 complex (Walther et al., 2003).

All Nup98 chromosomal translocations that have been observed have a breakpoint in the 3’ end of the Nup98 portion, disrupting the Nup98 open reading frame located upstream of Nup96 (Xu and Powers, 2009). Thus, Nup98 translocations result in fusions of the N-terminal region of Nup98 with the C-terminal region of a partner gene, which varies (Simon and Rout, 2014). This almost certainly disrupts the expression of Nup96 as well, which requires Nup98-dependent autocatalytic processing from the Nup98-96 precursor to be properly localized and functional (Fontoura et al., 1999; Rosenblum and Blobel, 1999).

While most of the attention on Nup98 translocations in cancer has focused on overexpressing the fusion partners, there is increasing evidence that the disruption of endogenous Nup98 and/or Nup96 contribute to cancer. For example, loss of one copy of Nup96 in the mouse leads to mildly enhanced proliferation of T-cells, supporting a potential role for Nup96 as a haploinsufficient tumor suppressor (Chakraborty et al., 2008), but Nup96+/- mice do not appear to exhibit cell cycle deregulation in other tissues nor develop cancer (Faria et al., 2006). Loss of one copy of Nup98 in the mouse (with Nup96 remaining intact) cooperates with loss of the nuclear export cofactor Rae1 to increase aneuploidy (Jeganathan et al., 2005), but Nup98+/- mice have not been reported to develop cancer nor to exhibit cell cycle de-regulation on their own (Wu et al., 2001). Studies of Nup98 and Nup96 homozygous mutants have been severely limited by the very early embryonic lethality caused the by loss of each Nup (Faria et al., 2006; Wu et al., 2001), and compound mutants have not been reported. Using a small interfering RNA (siRNA) knockdown approach to selectively target Nup98 in human cells, revealed a role for Nup98 in p53-dependent induction of the cdk inhibitor p21 in response to DNA damage, consistent with a tumor-suppressor function for Nup98 (Singer et al., 2012). Furthermore, work in *Drosophila* revealed an unexpected off-pore role for Nup98 in modulating the expression of several cell cycle genes (Capelson et al., 2010; Kalverda et al., 2010). Human *NUP98-96* is located near a known imprinted tumor-suppressor region in the genome (Joyce and Schofield, 1998), which could be significant as loss of heterozygosity via mutation or epigenetic modifications for the remaining Nup98-96 locus may occur in cancers exhibiting translocations. We are not aware of any information reported to date about the expression levels from the non-translocated *NUP98-96* gene in these diseases.

We simultaneously inhibited Nup98 and 96 in *Drosophila* using an *in vivo* RNAi knockdown approach and observed cell cycle de-regulation and cooperation with oncogenic mutations, consistent with a tumor suppressor function for Nup98 and/or 96. Transgenes encoding Nup98 or Nup96 individually do not rescue this phenotype, while expression of a transgene encoding both does – suggesting Nup98 and Nup96 play non-overlapping and potentially synergistic roles in cell cycle regulation.

Here we show that that reducing Nup98-96 function via an RNAi approach in *Drosophila melanogaster* (where the Nup98-96 shared mRNA and reading frame gene structure is conserved) de-regulates the cell cycle. We find evidence of overproliferation in Nup98-96 deficient tissues, counteracted by elevated apoptosis and aberrant Wingless and JNK signaling associated with wound healing. When the knockdown of Nup98-96 is combined with inhibition of apoptosis, we see synergism leading to overgrowth consistent with a tumor-suppressor function for endogenous Nup98 and/or 96. We suggest that the loss of normal Nup98 and Nup96 function de-regulates the cell cycle to cooperate with other mutations in cancer.

## Results

### Loss of Nup98-96 disrupts G1 arrests and causes cell cycle de-regulation

We previously described an RNAi screen to identify genes that promote proper cell cycle exit in the *Drosophila* eye (Flegel et al., 2016; Sun and Buttitta, 2015). Our initial screen used UAS-RNAi constructs from the Harvard TRiP RNAi collection, driven by the *Glass Multimer Repeats* (*GMR*) promoter-Gal4 with an E2F-responsive *PCNA-white* reporter transgene, which provides adult eye color as a readout of E2F and cell cycle activity (Bandura et al., 2013). This screen successfully identified genes that delay proper cell cycle exit by promoting a delay or bypass of G1 arrest, which directly or indirectly impacts E2F activity (Flegel et al., 2016; Sun and Buttitta, 2015). In this screen, we identified an RNAi line targeting the bi-cistronic Nup98-96 transcript as a potential novel regulator of cell cycle exit in the *Drosophila* eye.

Cell cycle exit in the eye is normally completed by 24 hours after puparium formation (APF). To confirm whether knockdown of Nup98-96 delayed cell cycle exit in the pupa eye, we performed S-phase labeling via EdU incorporation and examined an E2F transcriptional activity reporter *PCNA-GFP* in pupal eyes several hours after normal cell cycle exit. We confirmed that knockdown of Nup98-96 delayed proper cell cycle exit in the pupa eye to between 28-36h APF (Supp Fig. 1A). We also confirmed that the RNAi line identified in the screen knocked down endogenous Nup98-96 tagged with GFP and that re-expression of both exogenous Nup98 and Nup96 were required to rescue phenotypes due to Nup98-96 bi-cistronic transcript knockdown (Supp Fig. 1 B,C). Neither exogenous Nup98 or Nup96 alone were sufficient to rescue Nup98-96 RNAi phenotypes, suggesting both Nups contribute to the cell cycle exit defect.

We next examined whether knockdown of Nup98-96 in the posterior wing using the driver *engrailed-Gal4* (*en-Gal4*) with a temperature sensitive Gal80 (*en^TS^*) could delay cell cycle exit in the pupal wing, which also completes the final cell cycle by 24h APF. We used *Gal80^TS^* to limit expression of the RNAi to pupal stages to avoid developmental delays and lethality and an RNAi to the eye pigment gene *white* (*white^RNAi^*), which has no effect on cell cycle exit served as a negative control (Flegel et al., 2016). Labeling S-phases with EdU incorporation from 26-28h APF and mitoses using anti-phosphorylated Ser10-Histone H3 (PH3) antibody revealed that knockdown of Nup98-96 delayed cell cycle exit in the wing until 28-30h APF (Fig 1A-D).

**Fig. 1.**
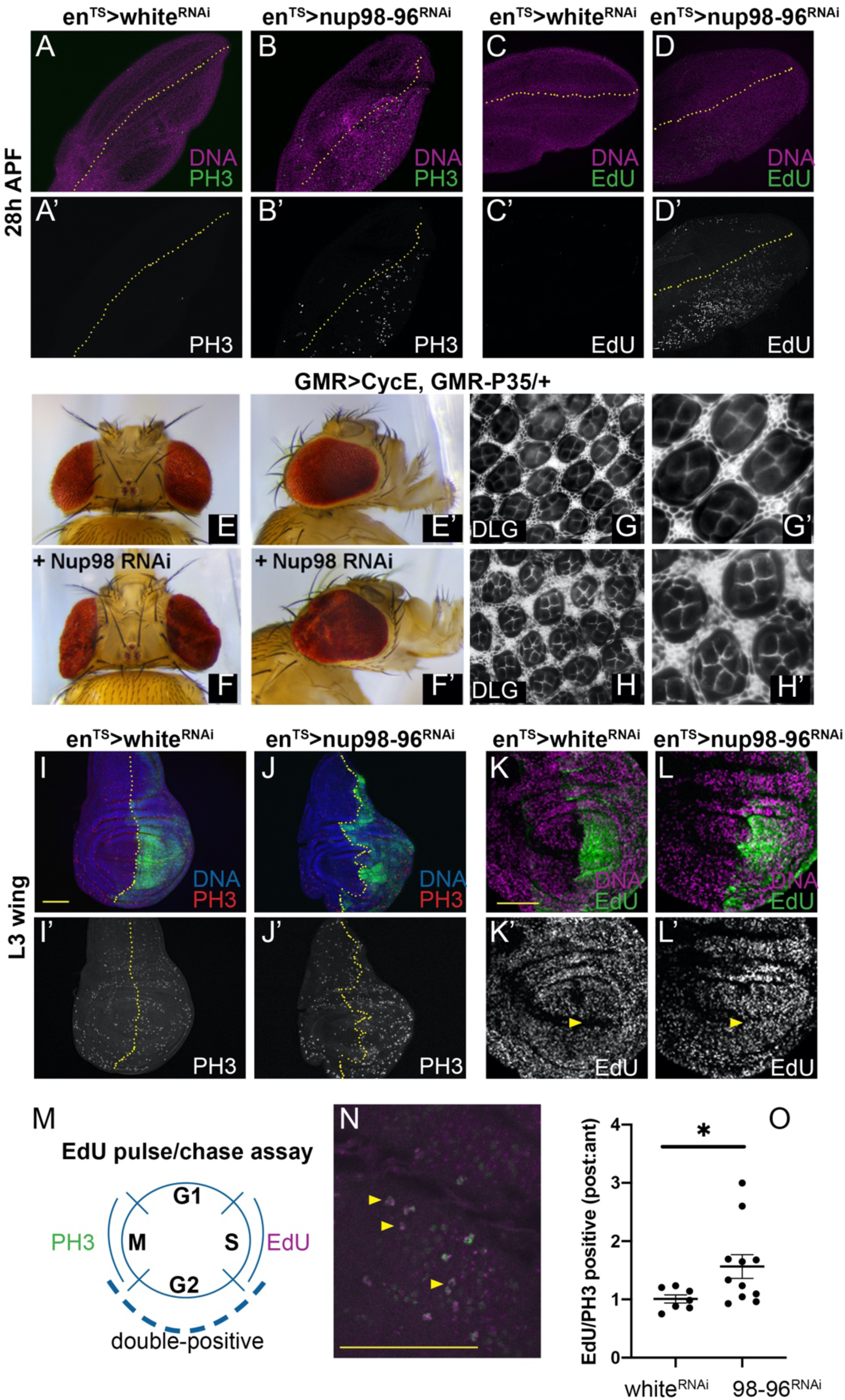
Inhibition of Nup98-96 leads to G1 bypass and cell cycle de-regulation. (A-D) Using *engrailed*-*Gal4* modified with a temperature sensitive Gal80 (*en*^TS^), the indicated UAS-RNAis were expressed in the posterior wing from mid L3 to 28h after puparium formation (APF) at 28°C. The dotted line indicates the pupal wing anterior/posterior boundary. Nup98-96 inhibition increased the number of mitoses (indicated by phospho-Ser10 histone H3, PH3) and S-phases indicated by 5-ethynyl-2-deoxyuridine (EdU) labeling in the posterior wing, at stages when the wing is normally post-mitotic. (E) Adult eyes from a heterozygous sensitized background expressing Cyclin E (CycE) under the GMR-Gal4 promoter and GMR-driven P35 is shown. (F) Adding in UAS-Nup98-96 RNAi enhanced eye size and folding, (G,H) and increased the number of cone cells and interommatidial cells as shown by staining for the septate junction protein Discs large (Dlg). (I-L) Using *en*^TS^, the indicated UAS-RNAis were expressed in the posterior wing for 72h prior to dissection of wandering L3. The dotted line indicates the anterior/posterior boundary. Nup98-96 inhibition increased the number of mitoses and S-phases in the posterior wing. A yellow arrowhead in K’-L’ indicates the posterior zone of non-proliferating cells (ZNC) which is normally G1 arrested, but undergoes S-phases when Nup98-96 is knocked down. (M-O) An EdU pulse for 1 h followed by a 7h chase and PH3 staining was used to label mid L3 wing cells that progress from S-M phase in ∼8h. (N) An example of a PH3 (green)/ EdU (magenta) double labeled disc is shown. (O) Quantification of double labeled cells in the posterior: anterior compartments normalizes for EdU incorporation in each wing and provides an indication of cell cycling speed differences between compartments. RNAi to Nup98-96 increased cycling speed in the posterior wing. (P<0.024; t-test with Welch’s correction). Yellow bar = 50 µm

We have shown that delays in cell cycle exit accompanied with high E2F activity can result from slowing the final cell cycle, or by causing additional cell cycles (Flegel et al., 2016; Sun and Buttitta, 2015). To determine which is the case with knockdown of Nup98-96, we expressed Nup98-96 RNAi in the eye, using a sensitized background with the GMR-Gal4 driver driving the G1-S Cyclin, Cyclin E (CycE) and the apoptosis inhibitor P35 (Hay et al., 1994). This sensitized background causes enlarged eyes and 1-3 extra cell cycles in the pupa eye prior to a robust cell cycle exit (Sun and Buttitta, 2015). The enlarged eyes of this sensitized background are visibly suppressed by factors that delay the cell cycle and enhanced by manipulations that cause extra cell cycles (Sun and Buttitta, 2015). Knockdown of Nup98-96 effectively enhanced the eye overgrowth of this sensitized background and resulted in extra cone cells and extra interommatidial cells in the pupal eye, confirming that the delay of cell cycle exit was caused by additional cell cycles (Fig. 1 E-H).

We next examined proliferating larval wings, to determine whether the effects of Nup98-96 knockdown were specific to the pupa or also impacted earlier cell cycles. We used *en-Gal4/Gal80^TS^* to express Nup98-96 RNAi in the posterior wing, labeled with GFP, for 72h prior to dissection and detected mitoses with PH3 or performed 5-10 min of EdU labeling for S-phase immediately prior to fixation. We observed an increase in mitoses when Nup98-96 was knocked down, accompanied by an increase in S-phase labeling (Fig. 1 I-L). Consistent with knockdown of Nup98-96 leading to a bypass of a G1 cell cycle arrest, we also observed abundant S-phases in the posterior zone of non-proliferating cells (ZNC, yellow arrowhead), which are normally quiescent at this stage (Johnston and Edgar, 1998). Similar effects on larval wing proliferation were observed using two independent Nup98-96 RNAi lines from the VDRC collection (Supp. Fig. 1D).

Increased EdU and PH3 labeling at fixed timepoints can only provide a snapshot of the cell cycle state in tissues but does not reveal cell cycle dynamics. To test whether the cell cycle is sped up when Nup98-96 is knocked down, we performed an EdU pulse/chase assay combined with PH3 labeling in L3 larval wings. We fed larvae with food containing EdU for 1 hour followed by a chase without EdU for 7h. At the end of the chase, we fixed larval wings and stained for PH3 and scored the number of cells double positive for EdU and PH3 in the posterior vs. anterior wing pouch for white RNAi vs. Nup98-96 RNAi wings. The posterior to anterior ratio of double-positive cells that transition from S to M-phase in control white RNAi wings is approximately 1, indicating similar cell cycle timing in the posterior and anterior wing pouch of late L3 larvae (Mesquita et al., 2010). By contrast, the ratio of double-positive cells was significantly increased when Nup98-96 was knocked down in the posterior, confirming that these cells also transition through the S/G2/M portion of the cell cycle faster (Fig. 1 M-O). An increased ratio could indicate both an increase in proliferation rate in the posterior wing, as well as a non-autonomous decrease in the anterior wing (Mesquita et al., 2010). The increased ratio of double-labeled cells in the Nup98-96 RNAi sample was not due to reduced labeling in the anterior compartment, as when we compared the total number of posterior double-labeled cells in Nup98-96 RNAi to posterior double-labeled cells in white RNAi wings (an external control), we observed a 20% increase in double-labeled cells when Nup98-96 was knocked down (Supp. Fig 1E).

### Nup98-96 knockdown results in apoptosis and activation of JNK signaling

Despite the increased rate of proliferation and disruption of G1 arrest in the larval and pupal tissues, we noted that the posterior wing expressing Nup98-96 RNAi was consistently smaller than normal (Supp. Fig, 1C). We therefore examined whether knockdown of Nup98-96 in the posterior larval wing increased cell loss via apoptosis. Knockdown of Nup98-96 for 72h dramatically increased apoptosis in the posterior wing, as measured by anti-cleaved Caspase 3 and anti-DCP1 staining (Fig. 2 A-B, Supp. Fig. 2). The increased apoptosis and reduced size in the posterior wing could be fully rescued by exogenous expression of both Nup98 and 96 in the presence of Nup98-96 RNAi (Supp. Fig. 1C, Supp. Fig 2C,D). Expression of Nup98-96 RNAi in the dorsal wing using *apterous-Gal4,Gal80^TS^* (*ap^TS^*) for 72h also induced robust apoptosis, indicating that the effect was not specific to the posterior wing (Supp. Fig. 2E). We knocked down the initiator caspase Dronc or effector caspase Drice in attempt to rescue the apoptotic cells, but neither fully suppressed the apoptotic response to Nup98-96 knockdown (Fig. 2 C,D), nor did co-expression of a dominant negative form of p53 (not shown, Brodsky et al., 2000). We next co-expressed the baculoviral caspase inhibitor P35 with Nup98-96 RNAi, which suppressed apoptosis (Supp Fig 2) and resulted in dramatic wing overgrowth phenotypes, including folding of the epithelium and duplication of wings (Fig. 2 E,F). The overgrowth and duplication of wing tissues was reminiscent of a phenotype observed during wing damage and regeneration when JNK signaling is activated (Perez-Garijo et al., 2009; Schuster and Smith-Bolton, 2015; Verghese and Su, 2017; Worley et al., 2018). We therefore examined whether Nup98-96 knockdown resulted in activation of JNK signaling by staining for phospho-JNK (Fig. 2G,H) and induction of the JNK signaling transcriptional target *puckered* (using a *puc-LacZ* expression reporter, Supp. Fig. 2I). Knockdown of Nup98-96 for 72h led to high levels of compartment-autonomous JNK signaling in the wing.

**Fig. 2.**
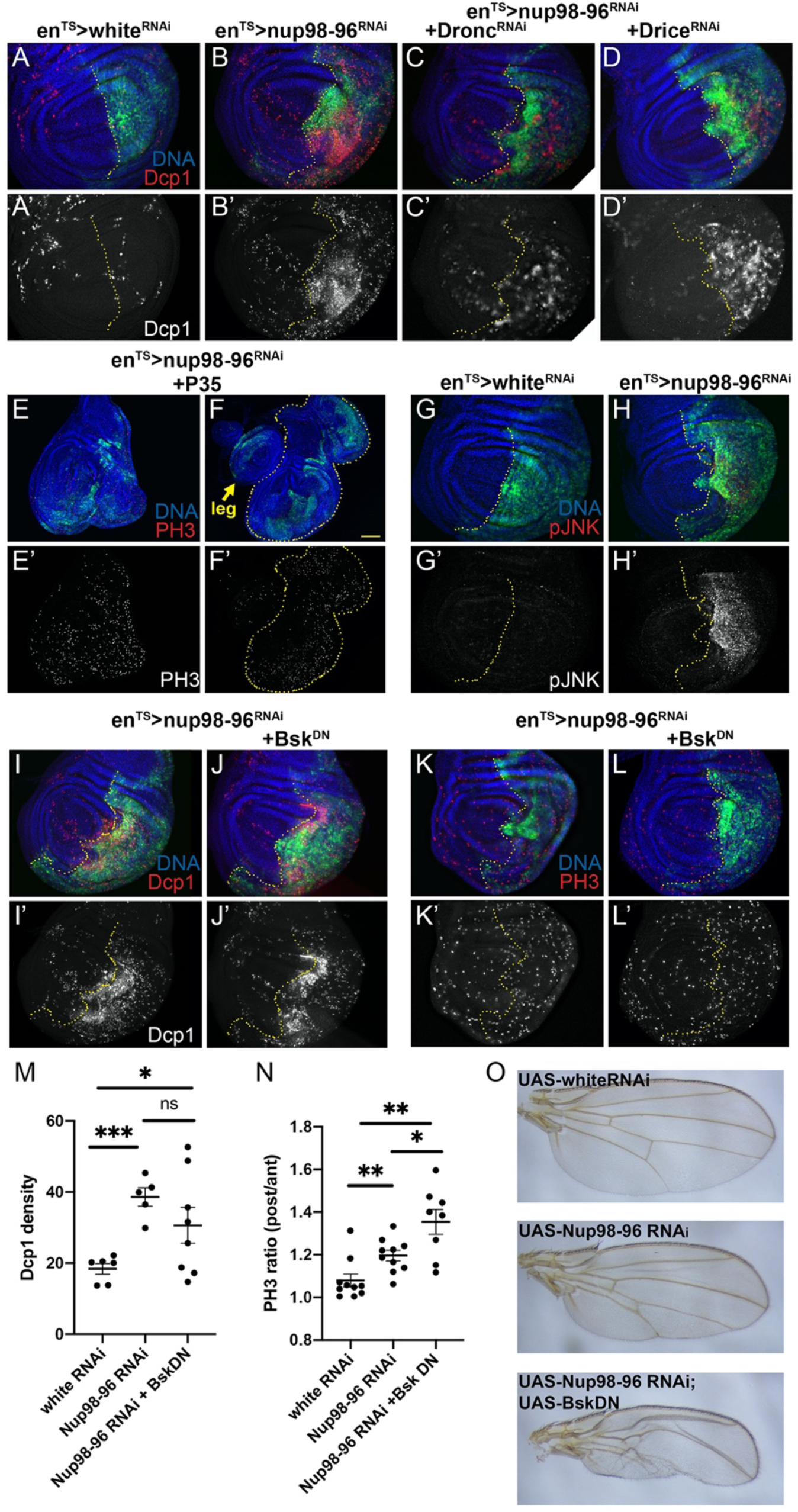
Inhibition of Nup98-96 leads to cell death and compensatory proliferation. (A-L) Using *en*^TS^, the indicated UAS-RNAis were expressed in the posterior wing for 72h prior to dissection of wandering L3 (unless otherwise indicated). The dotted line indicates the anterior/posterior boundary. (A-D) Nup98-96 inhibition increased apoptosis in the posterior wing, as indicated by cleaved Death Caspase-1 (Dcp1). (E-F) Co-expression of UAS-P35 with Nup98-96 RNAi lead to tissue overgrowth (E) and by day 5, wing pouch duplication (F). (G-H) Nup-98-96 knockdown led to activation of JNK signaling as detected by phosphorylated JNK staining (pJNK). (I-N) Co-expression of a dominant negative form of Drosophila JNK, Basket (Bsk^DN^) had variable effects on Dcp1 staining and increased the ratio of PH3 labeling in posterior:anterior wings, although overall PH3 signal decreased with Bsk^DN^ (Supp. Fig. 2). (Welch’s t-test comparisons, ns= not significant, *P<0.05, ** P<0.01, ***P<0.005.). Yellow bar = 100µm

High JNK signaling can paradoxically lead to both proliferation and cell death in *Drosophila* tissues (Fogarty and Bergmann, 2017). We next tested whether inhibition of JNK signaling via dominant negative form of the *Drosophila* JNK, Basket (Bsk^DN^) could suppress the apoptotic and proliferative response to knockdown of Nup98-96. Co-expression of Bsk^DN^ with Nup98-96 RNAi had a complex effect on apoptosis in the wing, enhancing levels of apoptosis in some samples, while suppressing in others (Fig. 2 I-J,M). Unexpectedly, co-expression of Bsk^DN^ with Nup98-96 RNAi did not suppress the increased mitoses observed in posterior wings expressing Nup98-96 RNAi, and even mildly enhanced the differences in mitotic labeling between anterior and posterior compartments (Fig. 2K-L, N). Although, we noted an overall decrease in PH3 labeling across both compartments when Bsk^DN^ was co-expressed in the posterior wing (Supp. Fig 2J), suggesting blocking JNK signaling reduced compensatory proliferation both autonomously and non-autonomously. The few adult wings that could be recovered with both Nup98-96 RNAi and Bsk^DN^ exhibited a more severely reduced posterior compartment than Nup98-96 RNAi alone (Fig. 2O). This suggests activation of JNK signaling provides compensatory proliferation and may partially increase survival when Nup98-96 is knocked down, consistent with previously described roles in wing damage and regeneration (Bergantinos et al., 2010; Herrera et al., 2013).

### Nup98-96 knockdown leads to mis-patterning and gene expression resembling a wound healing and loser phenotype

The JNK signaling and overgrowth phenotypes caused by suppressing apoptosis during Nup98-96 knockdown, are reminiscent of a phenomenon called apoptosis-induced compensatory proliferation (AIP) (Fogarty and Bergmann, 2017), which can impact tissue patterning. As previously described for other JNK-driven Drosophila tumor models, we observed dramatic tissue folding and invasion behaviors at both the A-P and D-V compartment boundaries when Nup98-96 was inhibited with P35 expression, (Supp. Fig 3A-C) (Muzzopappa et al., 2017). Therefore, we next investigated whether wing disc patterning is disrupted by Nup98-96 knockdown as previously shown in AIP.

We first examined Wg levels in wings expressing Nup98-96 RNAi, since AIP and wing duplications have been associated with ectopic Wg (Baonza et al., 2000; Perez-Garijo et al., 2009; Verghese and Su, 2017; Worley et al., 2018). We found that knockdown of Nup98-96 resulted in ectopic Wg in the dorsal wing hinge and this effect was amplified in in the presence of P35 (Fig. 3 A-D). We also observed ectopic phosphorylation of the transcription factor Mad (Supp. Fig. 3D), consistent with the previously described effect of AIP on Dpp signaling (Perez-Garijo et al., 2009; Pinal et al., 2018).

**Fig. 3.**
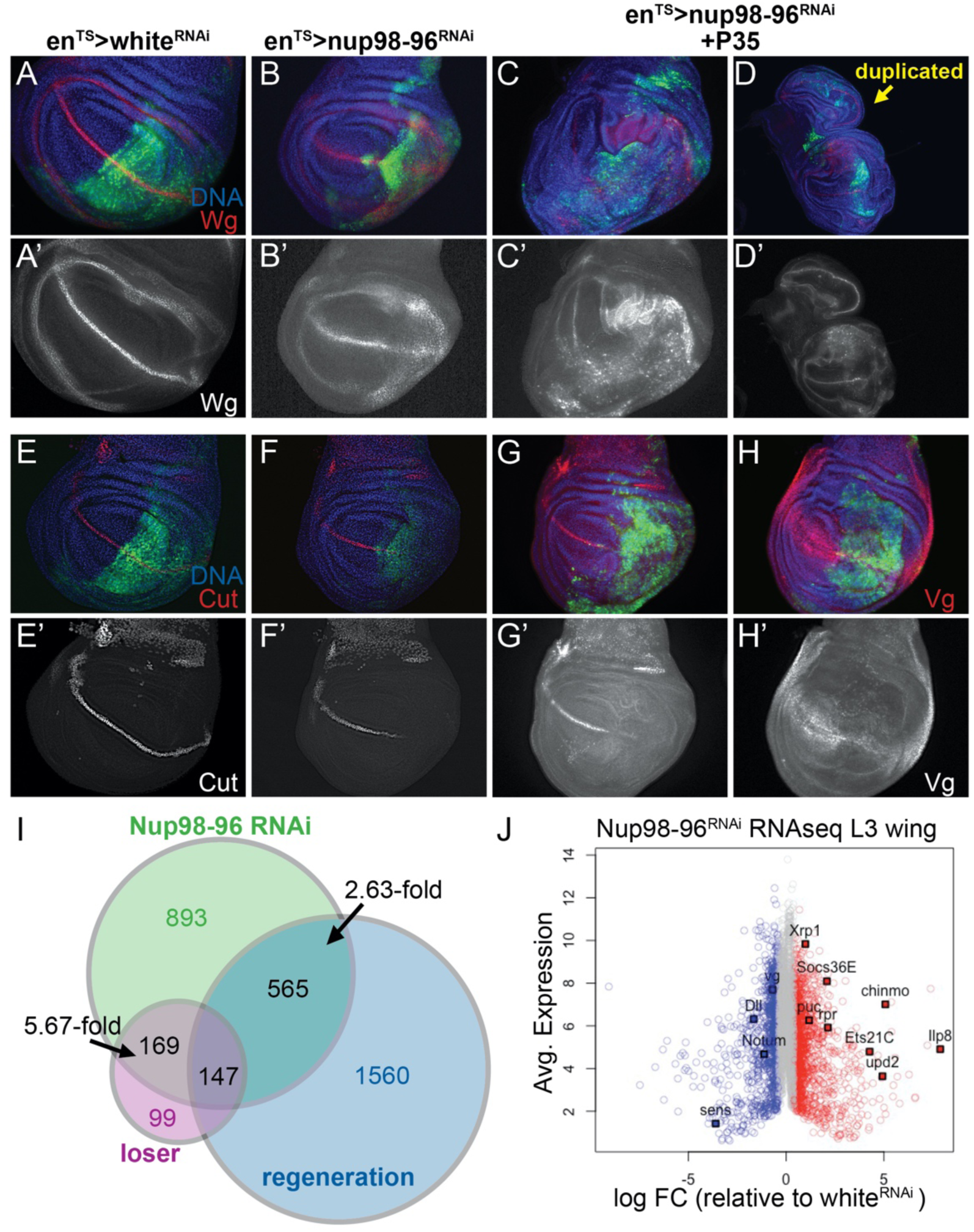
Inhibition of Nup98-96 leads to mis-patterning and gene expression changes associated with wounding and a “loser” phenotype. (A-H) Using *en*^TS^, the indicated UAS-RNAis were expressed in the posterior wing for 72h prior to dissection of wandering L3 (unless otherwise indicated). Discs in C, D, G and H co-express P35 to block apoptosis and allow for tissue overgrowth. Samples in C and D were dissected after 5 days of Nup98-96 RNAi+P35 expression. (A-D) Wg levels are disrupted at the Dorso-Ventral (DV) margin but increased at the dorsal hinge upon Nup98-96 knockdown. The effect on Wg and wing overgrowth is enhanced with P35. (E-G) Cut expression at the DV margin is disrupted by Nup98-96 knockdown, independent of P35 expression. (H) Vestigial (Vg) is reduced when Nup98-96 is knocked down. (I-J) RNAseq was performed on dissected late L3 wings expressing UAS-Nup98-96 or white RNAi for 72h, driven by *apterous-*Gal4 with *tub-Gal80^TS^ (ap^TS^)*. (I) A comparison of the overlap of genes significantly altered by Nup98-96 RNAi (1.5-fold or more) to previously published “wounding” and “loser” gene expression signatures in wings. The fold-enrichment in the overlap of genes, above that expected by chance is shown. (J) An M-A plot of the RNAseq data with significantly increased expression indicated in red, and significantly decreased expression in blue. Genes in grey are not significantly altered. Yellow bar = 100µm.

Both Wg and Notch have been implicated in the G1 arrest in the posterior ZNC (Duman-Scheel et al., 2004; Herranz et al., 2008). We therefore next examined the expression of two targets of Notch and Wg signaling; Cut, which is expressed in G1 arrested cells at the Dorso-Ventral (D-V) boundary, and Vestigial (Vg), which is expressed in a broader domain of the pouch induced by longer-range Wg signaling (de Celis et al., 1996; Kim et al., 1996; Neumann and Cohen, 1997). We found that Cut expression at the D-V boundary was nearly eliminated when Nup98-96 was knocked down, both with and without P35 (Fig. 3 E-G). This suggests Notch signaling at the D-V boundary is compromised when Nup98-96 function is reduced. Vg, an important wing identity and growth regulator (Halder et al., 1998; Williams et al., 1991; Williams et al., 1993; Zecca and Struhl, 2010), was also dramatically reduced in the pouch upon Nup98-96 knockdown (Fig.3H) suggesting Wg released from the D-V boundary is also compromised. Notch and Wg have been suggested to regulate the ZNC cell cycle arrest via repression of dMyc expression, but we did not observe any effects of Nup98-96 knockdown on dMyc levels in the ZNC (not shown). Interestingly, the downregulation of Vg was also observed in regenerating discs (Smith-Bolton et al., 2009), potentially due to the replacement of dying pouch cells with cells from the neighboring areas of the wing (Zecca and Struhl, 2010). Taken together, these data demonstrate that reduction of Nup98-96 function in the presence of P35 leads to AIP and wing mis-patterning and cell identity changes associated with a chronic wounding and regeneration response.

While high JNK signaling and apoptosis-induced compensatory proliferation can explain many of the phenotypes we observe with Nup98-96 knockdown, this does not reveal the proximal defect caused by loss of Nup98-96 function. To determine additional effects of Nup98-96 knockdown on gene expression in the wing, we performed comparative gene expression analysis via RNAseq to identify mRNAs increased or decreased upon Nup98-96 RNAi compared to the control white RNAi for 72h in late L3 wings (Supplemental Table 1). We observed the strong upregulation of many genes directly associated with JNK signaling (e.g. *puc, mmp1, Ets21C*) (Kulshammer et al., 2015; McEwen and Peifer, 2005; Uhlirova and Bohmann, 2006), Jak/STAT signaling (*upd, upd2, Socs36E*) (Amoyel et al., 2014) and developmental delays associated with wing damage and regeneration (*chinmo, Ilp8*) (Colombani et al., 2012; Garelli et al., 2012; Katsuyama et al., 2015; Narbonne-Reveau and Maurange, 2019). Consistent with increased cell cycle progression, we also observed the upregulation of several DNA damage and replication genes regulated by E2F activity (*Orc1*, multiple DNA Polymerases, *SpnE*, *Rnr-L*, *RfC4*) (Buttitta et al., 2010; Dimova et al., 2003). However, we did not observe strong upregulation of other G1-S promoting genes such as *dMyc* (1.52-fold change), *bantam*, *cycE* or *cycD*. When we compared gene expression signatures globally, we found a strong overlap (2.63- fold more genes than expected by chance) with a wounding and regeneration gene expression signature (Khan et al., 2017, Supplemental Table 2). We also noted upregulation of several genes associated with proteotoxic and oxidative stress (*Xrp1*, multiple Glutathione S transferases, *Aox1,* and specific DNA damage response genes) (Baumgartner et al., 2021). We found the strongest overlap of the Nup98-96 knockdown signature with a cell competition “loser” gene expression signature (5.67-fold more genes than expected by chance, 316/443 genes, Supplemental Table 3), which is also known to activate chronic JNK signaling (Kucinski et al., 2017).

### Nup98-96 knockdown leads to defects in proteins synthesis

The strong overlap of the gene expression changes in Nup98-96 knockdown with the cell competition “loser” signature suggested to us that a proximal effect of Nup98 loss could be on ribosome biogenesis. We further examined a gene expression signature associated with Xrp1, which mediates signaling downstream of ribosomal protein mutations (Lee et al., 2018). We found a striking proportion of Xrp1 targets (115 out of 159 overlapping in our dataset, Supplemental Table 4) were upregulated when Nup98-96 was knocked down (Ji et al., 2019). Consistent with a defect in ribosome function, we observed a decrease in protein synthesis when Nup98-96 was knocked down in wings, as measured by a puromycin labeling assay (Deliu et al., 2017), (Fig. 4 A,B). We did not observe downregulation of any ribosomal proteins in our RNAseq dataset, with the exception of a 2-fold decrease in *RpS19b*, which is a non-minute, duplicated ribosomal protein gene with tissue-specific expression (Marygold et al., 2007). Any effects on *RpS19b* levels are likely buffered by its paralog *RpS19a,* which exhibits much stronger expression in larval wings and was unchanged by Nup98-96 knockdown (Brown et al., 2014).

**Fig. 4.**
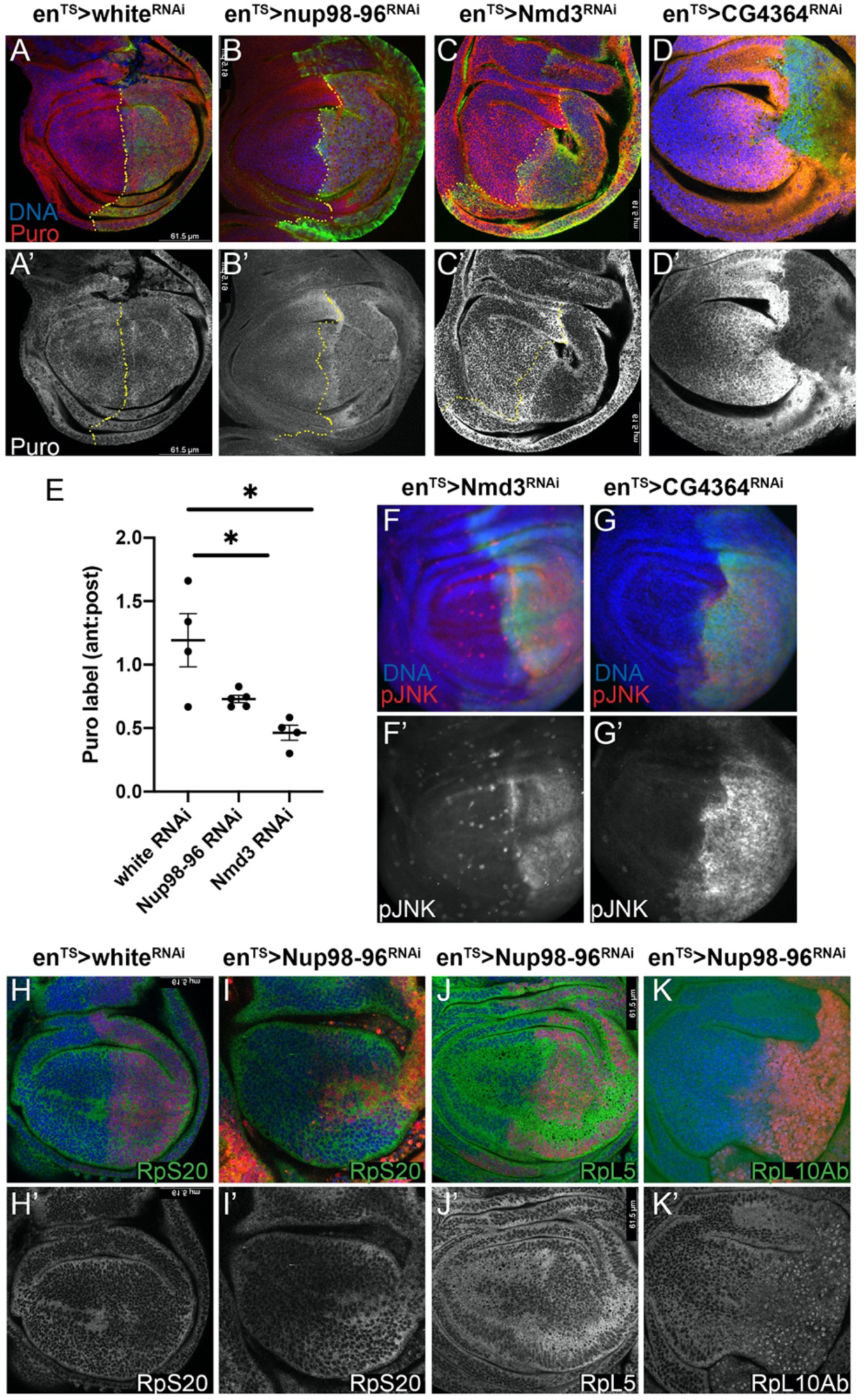
Knockdown of Nup98-96 leads to ribosomal protein mislocalization and compromised protein synthesis. (A-D) Using *en*^TS^, the indicated UAS-RNAis were expressed in the posterior wing for 72h prior to dissection of wandering L3 and labeled for protein synthesis using O-propargyl-puromycin (puro) incorporation. (E) The ratio of anterior:posterior puro-labeling is used to normalize for puro incorporation. Nup98-96 and Nmd3 knockdown reduced puro labeling (*P<0.05, unpaired t-test). (F-G) Knockdown of Nmd3 or CG4364 (Pescadillo homolog) for 48h in the posterior wing using *en^TS^* activated JNK signaling. (H-K) Using *enRFP*^TS^, the indicated UAS-RNAis were expressed for 72h in backgrounds expressing GFP or YFP-protein traps for the indicated Rp subunits. (K) RpL10Ab-YFP shows an aberrant nuclear enrichment when Nup98-96 is knocked down.

Nups play a key role in the CRM1-mediated export of ribosomal subunits (Gleizes et al., 2001; Johnson et al., 2002; Moy and Silver, 2002; Oeffinger et al., 2004), so we wondered if the proximal defect in Nup98-96 knockdown tissues might be defects in nuclear export of ribosomal complexes. First, we examined whether our partial knockdown of Nup98-96 function by RNAi was sufficient to disrupt nucleo-cytoplasmic localization, since previous work had suggested knockdown of Nup98-96 transcripts in *Drosophila* S2 cells did not produce such defects (Sabri et al., 2007). We confirmed that by 52h of knockdown with *en^TS^ in vivo*, we could easily visualize defects in nuclear localization of a ubiquitously expressed RFP with a nuclear localization signal (NLS). By 72h of knockdown, nuclear localization of NLS-RFP was almost completely abolished (Supp. Fig 4 A). We next confirmed that knockdown of an essential component of the nuclear export machinery for ribosome subunits, Nmd3 (Ma et al., 2017) also effectively reduced protein synthesis (Fig. 4 C). As a positive control we also knocked down CG4364, the fly homolog of the pre-rRNA processing component Pescadillo (Lapik et al., 2004) (Fig. 4 D-E). Inhibition of ribosome export machinery and pre-rRNA processing were both sufficient to induce strong JNK signaling (Fig. 4 F-G) in the wing.

Ribosome large and small complexes are exported from the nucleus separately as assembled pre-ribosomal particles and must associate with cytoplasmic maturation factors to exchange specific components to form mature functional ribosomes (Lo et al., 2010). We screened through collections of endogenously tagged Rp subunits and found RpL10Ab, but not other Rp subunits (RpS20 and RpL5) were mis-localized when Nup98-96 was knocked down (Fig. 4 H-K). Interestingly, the defect in RpL10Ab localization was nuclear retention, the opposite of the effect of Nup98-96 knockdown on NLS-RFP. RpL10Ab (also called L10a or uL1) is required to associate with Nmd3 for efficient pre-60S nuclear export (Musalgaonkar et al., 2019). Normally, RpL10Ab is translated in cytoplasm, localized to the nucleolus for assembly into the pre-60S complex and then exported bound to the Nmd3 adaptor. The nuclear retention of RpL10Ab upon Nup98-96 knockdown was initially puzzling as the other RpL subunits examined did not exhibit similar localization defects. However recent work has revealed that in mammals RpL10A is associated with a subset of specialized ribosomes and is not found in all 60S complexes (Shi et al., 2017). We suggest that knockdown of Nup98-96 partially compromises protein synthesis by inhibiting proper cytoplasmic translocation of a subset of pre- 60S subunits that are RpL10Ab-associated. Importantly, RpL10Ab is not a Minute gene (Marygold et al., 2007), possibly because it is a sub-stoichiometric ribosome component. Consistent with this, we do not recover significant overlap with the proteasomal stress portion of the “Loser” gene expression signature when Nup98-96 is compromised (Baumgartner et al., 2021).

### Nup98-96 knockdown in mammalian cells leads to defects in proteins synthesis and JNK activation

As described in the introduction, there is abundant evidence that loss of Nup98-96 function might contribute to tumorigenesis. We wondered whether inhibition of Nup98-96 in mammalian cells would also impact protein synthesis and JNK signaling as we observe in *Drosophila*. Of note, a screen for factors involved in ribosome biogenesis in HeLa cells identified several Nups containing FG repeats, including Nup98 as hits involved in pre-60S export, suggesting Nup98 effects on protein synthesis will be broadly conserved (Wild et al., 2010). We used small-interfering RNA (siRNA) to Nup98-96 in MCF7 breast cancer cells for 72 hours and compared effects on Nup98 levels, protein synthesis and pJNK to a control scrambled siRNA (ctrl siRNA). We found that the siRNA to Nup98-96 only partially reduced Nup98 levels, but that this reduction was sufficient to reduce protein synthesis and increase pJNK levels (Fig. 5 A-H).

**Fig. 5.**
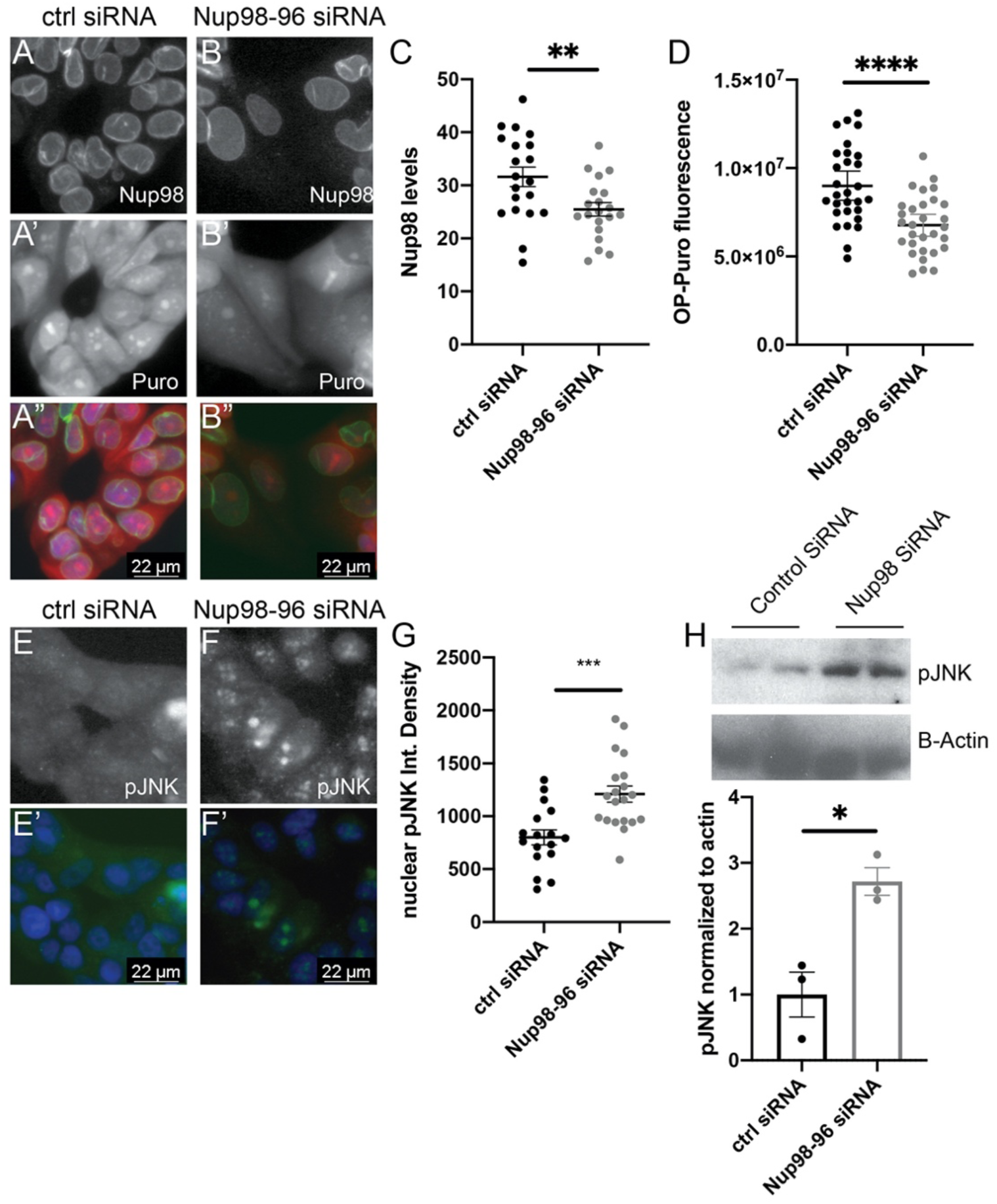
Knockdown of Nup98-96 in mammalian cells leads to reduced protein synthesis and JNK signaling. (A-B, E-F) MCF7 cells were treated with small interfering (si) RNAs for 72h and cells were labeled with Puro and fixed and stained with anti-Nup98 antibody (A-B), or fixed and stained for phospho-JNK (E-F). Control siRNA (Ctrl) is a scrambled siRNA. Nup98 siRNA reduces Nup98 levels (C) as well as reduced protein synthesis (D) and increases pJNK labeling (G-H). Quantifications of fluorescence were performed on individual cells from at least two experiments. ****P<0.0001, ***P<0.001, **P<0.01, *P<0.05 by unpaired t-tests, (G) uses Welch’s correction for unequal sample size.

### Overexpression of Nup98 leads to defects in proteins synthesis and JNK activation

Most of the attention on Nup98 translocations in cancer has focused on overexpressing Nup98 fusion partners. However when overexpressed, Nup98 has been shown to behave as a dominant negative and disrupt the nuclear envelope and nuclear transport (Fahrenkrog et al., 2016; Mendes et al., 2020), possibly by forming phase-separated aggregates outside of the nuclear pore (Ahn et al., 2021; Schmidt and Gorlich, 2015). We noted that Nup98 overexpression in the posterior wing reduced tissue size, and in severe cases disrupted pattering (Fig. 6 A-F). We therefore examined whether Nup98 overexpression in the *Drosophila* wing mimicked aspects of Nup98-96 inhibition. Overexpression of a strong UAS-Nup98 cDNA construct (2F) disrupted nuclear localization of an NLS-tagged RFP resulting in cytoplasmic accumulation (Fig. 6 G-H). Overexpression of a UAS-Nup98-96 cDNA construct was also sufficient to increase cell death and activate JNK signaling in the posterior wing (Fig. 6 I-J), and overexpression of both UAS- Nup98 and 96 or UAS-Nup98 alone (2F) reduced protein synthesis levels (Fig. 6 K-M). We suggest that Nup98-96 acts as a “goldilocks” gene (Braune and Lendahl, 2016), where too much or too little activity leads to stress signaling and acquisition of hallmarks of tumorigenesis. This complication makes this locus particularly prone to mis-regulation by translocations that reduce Nup98-96 normal functions and simultaneously provide additional Nup98-containing fusion proteins.

**Fig. 6.**
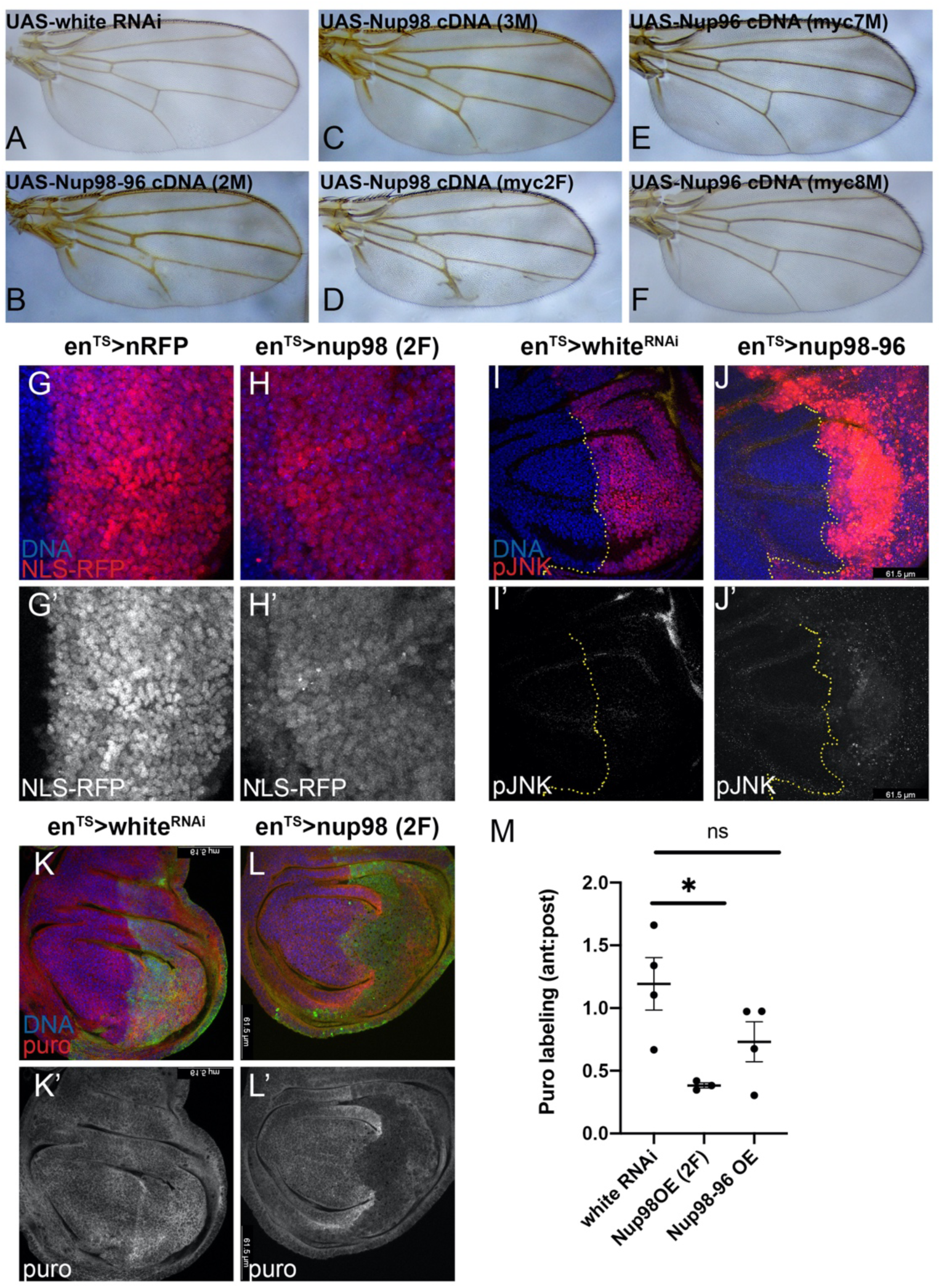
Overexpression of Nup98 disrupts protein synthesis and activates JNK signaling. (A-F) Using *en*^TS^, the indicated UAS-cDNA constructs were expressed in the posterior wing from mid-L2 and adult wings were mounted. Overexpression of Nup96 had no effect on the posterior wing while overexpression of Nup98 or Nup98-96 reduced posterior wing size and disrupted vein patterning. (G-H) Using *en*^TS^ RFP-NLS was expressed for 72h alone or together with UAS-Nup98 2F. Overexpression of Nup98 led to cytoplasmic accumulation of RFP, detected by the loss of discrete nuclear labeling. (I-J) Using *en*^TS^, Nup98-96 cDNA was expressed in the posterior wing for 72h prior to dissection of wandering L3 and labeling with pJNK. UAS-white RNAi serves as a negative control showing endogenous pJNK at this stage is very low. (K-M) Using *en*^TS^, the indicated UAS-cDNA or RNAi was expressed for 72h prior to dissection and labeling with puro to measure protein synthesis. Overexpression of Nup98 2F reduced protein synthesis in the posterior wing, which Nup98-96 overexpression had a milder effect. (*P<0.05 by unpaired t-test).

## Discussion

### Partial Nup98-96 loss of function leads to paradoxical increases in cell cycling and cell death accompanied by reduced protein synthesis

Protein synthesis and the cell cycle are usually coupled by pathways such as insulin and TOR signaling as well as growth and cell cycle checkpoints, which promote or limit cell cycle progression and protein synthesis coordinately (Grewal, 2009; Lockhead et al., 2020; Romero-Pozuelo et al., 2017; Romero-Pozuelo et al., 2020). Here, we describe a seemingly paradoxical situation where protein synthesis and cell cycle are effectively uncoupled. When Nups 98 and 96 are partially compromised, cells with reduced protein synthesis cycle more and even bypass developmentally induced G1 arrests. This is accompanied by high levels of chronic JNK signaling and induction of apoptosis, along with expression of genes involved in tissue regeneration and compensatory proliferation. When apoptosis is blocked using the caspase inhibitor P35, tissue overgrowth and mis-patterning results, reminiscent of tumorigenesis. We propose that mutations or gene expression changes that reduce Nup98 and Nup96 function, in the presence of apoptosis suppression, can contribute to tumorigenesis. This may help explain contexts of Nup98 and/or Nup96 loss that could pre-dispose for cancer (Franks and Hetzer, 2013; Simon and Rout, 2014; Singer et al., 2012).

The phenotype we describe here for Nup98-96 inhibition is strikingly similar to that recently described for a ribosomal protein mutant, when cell death is blocked (Akai et al., 2021). When we examined the gene expression signature in response to reduced Nup98-96, we observed a strong overlap with conditions of reduced protein synthesis caused by stoichiometric imbalances in ribosomal proteins (Kucinski et al., 2017; Lee et al., 2018). We suggest this effect of Nup98-96 inhibition is due to defects in nucleo-cytoplasmic transport of RpL10A, although we cannot rule out that localization of other ribosomal proteins may also be affected. Because the defect is in RpL10A localization, rather than levels, we were unable to rescue the Nup98-96 knockdown phenotypes with RpL10A overexpression. On the contrary, we observed several stress signaling phenotypes when we overexpressed RpL10A itself even in a wild-type background suggesting RpL10A levels must also be carefully controlled (Chaichanit et al., 2018; Wonglapsuwan et al., 2011). This may be of broader consequence to the *Drosophila* research community since Gal4/UAS-driven overexpression of this ribosomal protein is used for translatome profiling through translating ribosome affinity purification (Thomas et al., 2012). Importantly, localization of 40S and 60S subunits are not globally disrupted in our Nup98-96 knockdown conditions and protein synthesis is only partially reduced. We suggest that this is because RpL10A is a sub-stoichiometric component of ribosomes and that only the subset of ribosomes containing RpL10A are affected. In mammals RpL10A-containing ribosomes have been shown to translate genes required for cell survival and are depleted of those required for cell death (Shi et al., 2017). Whether this is the case for *Drosophila* RpL10A- containing ribosomes remains to be determined, although increasing RpL10A expression in *Drosophila* has been shown to affect E-cadherin and Insulin-like receptor levels, suggesting components of these pathways could be regulated by RpL10A levels (Chaichanit et al., 2018).

The effects of reducing Nup98-96 expression are likely to be pleiotropic, and we cannot rule out the possibility that Nup98 and 96 mis-regulation may also lead to more direct effects on the cell cycle, independent of JNK signaling and reduced protein synthesis. Indeed, when JNK signaling is blocked by a dominant negative, overall compensatory proliferation is significantly reduced, but Nup98-96 reduced tissue still exhibits a slightly higher mitotic index than tissue with normal Nup98-96 levels. This could be in part the result of a known Nup98 interaction with the APC/C which leads to aneuploidy when Nup98 levels are reduced (Jeganathan et al., 2006; Jeganathan et al., 2005). This interaction with the APC/C may also explain the disruption of terminal cell cycle arrest caused by reduced Nup98-96, as high APC/C activity promotes proper timing of the final cell cycle (Buttitta et al., 2010; Reber et al., 2006; Ruggiero et al., 2012; Tanaka-Matakatsu et al., 2007). We tested for aneuploidy using flow cytometry on wings and did not observe obvious accumulation of aneuploidy when Nup98-96 is knocked down, either with or without apoptosis inhibition. Alternatively, effects on nuclear export of cell cycle factors or their mRNAs may also contribute to the cell cycle phenotypes (Chakraborty et al., 2008), although we did not find obvious changes in protein levels or dynamics of Cyclins A or B. We also examined whether mis-regulation of transcriptional targets of Nup98 regulated through off-pore roles may explain the phenotypes we observe, but we did not find significant overlap of genes altered in our Nup98-96 knockdown with Nup98 targets determined by ChIP-seq in larval brains (Pascual-Garcia et al., 2017) or RNAseq in S2 cells (Kalverda et al., 2010). We found a mild enrichment (1.43-fold over that expected by chance) in the overlap of genes altered in our Nup98-96 knockdown with Nup98-ChIP seq targets in S2 cells. (Pascual-Garcia et al., 2017, Supplemental Table 5). Overall, the previously described wounding/regeneration and “loser” gene expression programs explain nearly half (49.7%) of the gene expression changes we observe in wings when Nup98-96 is reduced (Fig. 3).

### AIP and hallmarks of tumorigenesis in Nup98 cancers

Blocking apoptosis in cells with inhibited Nup98-96 leads to phenotypes consistent with sustained apoptosis-induced proliferation (AIP), which is thought to contribute to tumorigenesis in epithelia (Fogarty and Bergmann, 2017). Epithelial tumors exhibit wounding phenotypes, chronic inflammation and cell death (Dvorak, 1986; Karin and Clevers, 2016). Chronic AIP leads to sustained proliferation and results in abnormal, hyperplastic overgrowth (Perez-Garijo et al., 2009; Pinal et al., 2018). AIP therefore could contribute to overproliferation in epithelial cancers with disrupted Nup98-96 expression. But how might this relate to aberrant signaling in hematological malignancies related to Nup98 mis-expression? It is possible effects of Nup98 mis-regulation impact different tissue types through similar pathways, but with distinct outcomes. For example, expression of a NUP98-HOXA9 fusion in a *Drosophila* model with a normal Nup98-96 locus leads to hyperplastic over-proliferation in hematopoietic tissues but minimal effects in epithelial tissues (Baril et al., 2017), while loss of Nup98-96 in larval hematopoietic tissues leads to a loss of progenitors, a phenotype also observed upon inhibition of the ribosomal protein RpS8 (Mondal et al., 2014). Thus Nup98-96 loss likely has distinct yet overlapping effects in different tissue types. Nup98 mutations in leukemias are associated with mutations affecting apoptosis such as BCR-ABL, NRAS, or KRAS and ICSBP (Gabriele et al., 1999; Gough et al., 2011; Gurevich et al., 2006; Hu et al., 2016; Slape et al., 2008). Mouse models with Nup98 fusions exhibit increased apoptosis (Choi et al., 2008; Lin et al., 2005), and a zebrafish model of NUP98-HOXA9-driven leukemia upregulates Bcl2 to suppress apoptosis (Forrester et al., 2011). In a mouse model of Nup98-HoxD13-driven leukemia, loss of p300 leads to reduced apoptosis and enhanced activation of Jak/Stat signaling, reminiscent of signaling effects we see in AIP (Cheng et al., 2017). In our Nup98-96 RNAi experiments we reduce Nup98 protein levels to about 50-70% of the normal level, consistent with other studies using this RNAi approach (Pascual-Garcia et al., 2014). Our data suggests that this locus can behave as a dominant negative when the Nup98 portion is overexpressed through translocations as well as a haplo-insufficient tumor suppressor in some contexts. We propose that disruption of the Nup98-96 locus in cancers with or without Nup98-translocations may contribute to tumorigenesis through aberrant JNK signaling and AIP, in the presence of additional hits that block cell death.

## Acknowledgements

We thank Drs. Sofia Merajver, Catherine Collins, Helena Richardson and Maya Capelson for sharing flies and reagents. We thank the Bloomington (BDSC), Vienna (VDRC) and Kyoto (DGRC) *Drosophila* stock centers for providing stocks critical to this work. We also thank A. Sustar and the lab of Dr. Gerold Schubiger for sharing Ptc and Vg antibodies originally obtained from the S. Carroll and T. Kornberg Labs. This work in the Buttitta Lab was supported by The American Cancer Society (RSG-15-161-01-DDC), the University of Michigan Rogel Cancer Center Discovery Fund and the NIH (R01GM127367). We thank the U. Michigan Advanced Genomics Core for library preparation and high-throughput sequencing. We thank the Buttitta Lab members for helpful input on this project. L.A.B. thanks Lynn Taylor for essential childcare support during the writing of this paper.

## Methods

### Flystocks used

UAS-Nup98-96 RNAi (TRiP BL28562, VDRC lines, KK100388 and GD6897)

UAS-white RNAi (TRiP BL35573)

UAS-Dronc RNAi (TRiP BL 32963)

UAS-Drice RNAi (TRiP BL 32403)

UAS-Nmd3 RNAi (VDRC105619 and VDRC46166)

UAS-CG4364 RNAi (VDRC27607)

GMR-Gal4, UAS-CycE(I); GMR-P35 from H. Richardson

Nup98-96-GFP (VDRC 318656 FlyFos collection)

UAS-Nup98-96 cDNA(2M), UAS-Nup98 cDNA (3M), UAS-Nup98 cDNA (myc2F), UAS-Nup96 cDNA (myc7M), UAS-Nup96 cDNA (myc8M) all from M. Capelson. *En^TS^* is w; *en*-Gal4,UAS-GFP; *tub*-Gal80TS/TM6B from (Buttitta et al., 2007) *Ap^TS^* is w; *ap*-Gal4,UAS-GFP; *tub*-Gal80TS/TM6B from (Buttitta et al., 2007)

*En^TS^* RFP is w; *en*-Gal4,UAS-RFP_NLS_; *tub*-Gal80TS/TM6B

UAS-Bsk^DN^ (on III mutated in kinase domain) and *puc^e69^-LacZ* provided by C. Collins.

UAS-P35 on X (BL6298)

RpS20-GFP (Kyoto 109696 w^1118^; PBacRpS20^KM0175^ / TM2)

RpL5-GFP (Kyoto 109767 w^1118^; PBac RpL5^KM0174^ / SM6a and Kyoto 109768 w^1118^; PBac _RpL5_^KM0163^_)_

RpL10Ab-YFP (Kyoto 115462 w^1118^; PBac RpL10Ab^CPTI003957^)

*Ubi*-RFP_NLS_: (derived from BL35496)

*y*,*w*,*hs-flp12* (derived from BL1929)

*w*; *act*>stop>Gal4, UAS-GFP_NLS_; UAS-P35 from (Neufeld et al., 1998)

### Immunofluorescence

*Drosophila* samples were fixed in 4% paraformaldehyde/1XPBS solution for 20-30 min., rinsed twice in 1X PBS with 0.1% Triton-X-100 detergent (1XPBST). The samples were then incubated in appropriate dilution of antibodies in PAT (1XPBS + 0.1% Triton X-100 + 1% BSA) for 4 h at room temperature or overnight at 4°C. The samples were then washed thrice for 10 mins in 1XPBST and incubated in secondary antibody conjugated with required fluorophore for 4 h in PBT-X + 2% normal goat serum (1XPBS + 0.3% Triton X-100 + 0.1% BSA) at room temperature or overnight at 4°C. DAPI or Hoechsts 33258 was used as a nuclear counter-stain and samples were mounted on glass slides using 5µl of Vectashield mounting medium (Vector Labs). Slides were imaged using a Leica DMI6000 epifluorescence system with subsequent deconvolution or a Leica SP5 confocal microscope.

For MCF-7 cells, fixation and washes were performed as described above, except in 6- well dishes, with just 1h of incubation with primary and secondary antibodies at room tempertaure.

### EdU labeling and pulse-chase assay

Crosses were flipped every day and kept at room temperature (22°C). The vials with embryos were transferred to 29°C after 2 days. Larvae at mid-L3, (∼66 hrs after the transfer) were removed from the vials by floating in 30% Sucrose/1XPBS solution. The larvae were transferred to a vial with YG food mixed with 100uM EdU and blue food coloring (to track feeding) at 29°C for 1h. Larvae with blue abdomens were then transferred to fresh non-EdU food (chase) for 6h at 29°C (equivalent to 7h at 25°C). EdU pulsed-chased wandering L3 larvae were collected, dissected, fixed, and antibody stained for EdU, PH3 and GFP (to mark the anterior-posterior compartment boundary). The EdU labelling was performed using a Click it EdU-555 kit (Cat No C10338, Invitrogen) following the manufacturer’s instruction. The slide was then imaged using confocal microscopy and the total number of cells positive for both EdU and PH3 were scored.

### Protein synthesis puromycin assay

L3 larvae were dissected in Ringer’s solution (Sullivan, 2000) and inverted larvae heads containing wing discs were incubated with 20µm of OPP (O-Propargyl-Puromycin, Invitrogen) in Ringer’s solution for 12 mins. The sample was then fixed with 4% paraformaldehyde/1XPBS solution for 20-30 and labelled using the Click-it OPP kit (Cat No C10457, Invitrogen) following the manufacturer’s instruction.

### Antibodies used

Mouse anti-PH3 Cell Signaling 9707 1:1000

Rabbit anti-PH3 Millipore 06-570 1:2000

Rabbit anti-Dcp1 Cell Signaling 9578 1:100

Rabbit anti-pJNK Promega v7931 1:100 (for *Drosophila*, used slightly younger pre-wandering larvae due to high peripodial signal in later larvae)

Rabbit anti-pSmad Cell Signaling 9516 1:50 (dissection must be performed on ice)

Rabbit anti-GFP Invitrogen A11122 1:1000 (for co-labeling GFP with EdU)

Mouse anti-cut DSHB 2B10 1:100

Mouse anti-lamin Dm0 DSHB ADL67.10 1:100

Mouse anti-Wg DSHB 4D4 1:100

Rabbit anti-Vg (1:200) via G. Schubiger, from S. Carroll

mouse anti-Patched (1:200) via G. Schubiger, from T. Kornberg

### siRNA in MCF7 cells

MCF7 cells were a gift from the Merajver lab (U. Michigan). The cells were gown to 50-70% confluency in a 6-well plate. The cells were then transfected with 20nM of Nup98 SiRNA or control SiRNA using PEI max. The media was changed the next day and incubated for 72 hrs. The cells were then harvested for fixation and staining or lysed for western blot.

siRNAs: MCF7 Silencer™ Select Negative Control No. 1 siRNA Catalog number: 4390843

### Image analysis and quantification

Image quantification was performed using FIJI. For quantification of Dcp1 or pJNK labeling regions of similar size (ROIs) in the anterior and posterior wing were hand-drawn using the nuclear (Dapi or Hoeschsts 33258) staining to indicate tissue boundaries and GFP labeling for compartment boundaries. Area-normalized integrated density was used for Dcp1, PH3, Nup98 and puromycin quantification. For ratios in the EdU/PH3 pulse chase assay, double-labeled cells were counted in each compartment and the ratio across wings is shown. Each dot in the scatter plot represents and individual wing (For Figs 1,2, 4) or cells (Fig 5).

### Mounting and imaging of adult wings

Adult wings were preserved in Ethanol, washed in Methyl salicylate and mounted in Canada Balsam (Sigma) as described (O’Keefe et al., 2012). Adult wings were photographed under brightfield conditions on a Leitz Orthoplan2 at 5× magnification, using a Nikon DS-Vi1 color camera and Nikon NIS Elements software.

### RNAseq

Experimental animals contained the genotype: UAS-P35/w; ap-Gal4, UAS-GFP/ +; tub- gal80TS/UAS-Nup98-96 RNAi TRiP

Control animals contained the genotype: UAS-P35/w; ap-Gal4, UAS-GFP/ +; tub- gal80TS/UAS-white RNAi TRiP

Crosses were performed at room temperature and embryos were collected within a 12h window to synchronize developmental staging and shifted to 18°C. Animals were reared in uncrowded conditions (70 larvae per vial). On day 4 animals were transferred to 28°C and 72h later 3^rd^ instar wing discs were dissected in sterile 1X PBS. We followed a Trizol-based RNA preparation protocol with dounce homogenization of 40 wings per sample with 3 replicated per genotype, as previously described (Flegel et al., 2016).

Using PolyA selection, the University of Michigan’s Sequencing Core generated barcoded libraries for each sample and confirmed the quality via the Bioanalyzer and qPCR. Sequencing was performed with the Illumina HiSeq 2000 platform and high read quality was confirmed using FastQC. Reads were aligned to the BDGP6.82 D. melanogaster genome using Rsubread (v1.21.5), with featureCounts resulting in >77% of the reads being successfully assigned to genes (Liao et al., 2014). Counts per million (cpm) were determined with edgeR (v3.13.4) and transcripts with low expression were identified and removed using the data-based Jaccard similarity index determined with HTSFilter (v1.11.0). The cpm were TMM normalized (calcNormFactors), voom transformed (Law et al., 2014), fit to a linear model (lmFit), then differential gene expression calls were made with eBayes. The full dataset is available on GEO (GSE152679). For significance of overlap with other datasets (Figure 3), hypergeometric probabilities were calculated using the hypergeometric distribution as described (Flegel et al., 2016).

---

**Supp Fig 1:**
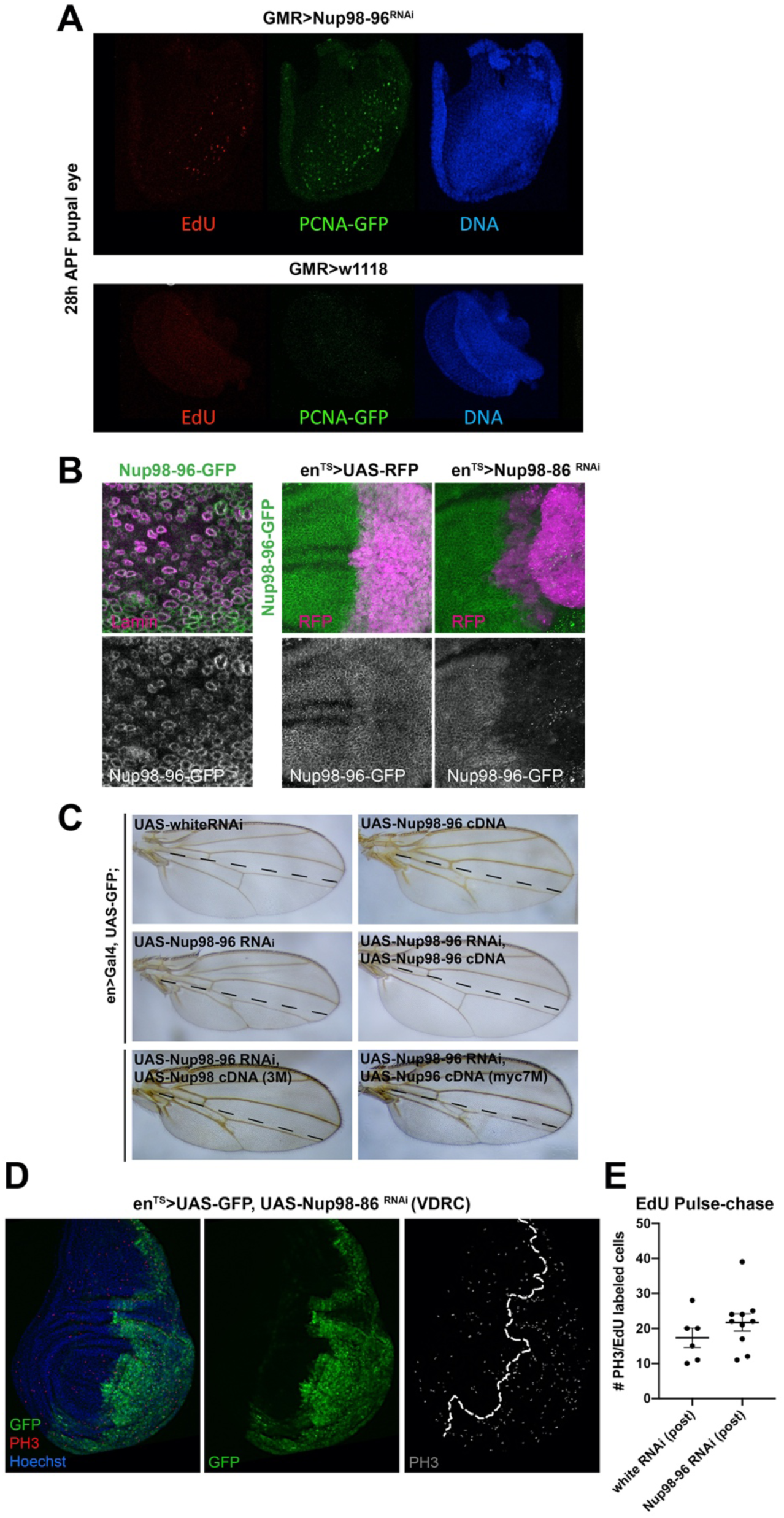
(A) 28h APF pupal eyes were labeled for 2h with EdU to detect S-phase and stained for GFP to detect PCNA promoter driven GFP. S-phases and PCNA promoter activity is evident when RNAi to Nup98-96 is driven with GMR-Gal4. GMR-Gal4 without any RNAi (in a w1118 background) serves as a control. Genotypes are: (top) w; GMR-Gal4 /+; UAS-Nup98-96 RNAi (TRiP)/PCNA prom-GFP (bottom) w; GMR-Gal4/+; PCNA prom-GFP/+. (B) Nup98-96-GFP is from the FlyFos collection containing a Fosmid on II with Nup98-96 coding region plus regulatory DNA and a GFP tag (Sarov et al., 2016) We confirmed this line exhibits the expected ubiquitous nuclear envelop labeling by co-staining with Lamin Dm0. (Right) We co-expressed Nup98-96-GFP in a background with en-Gal4 driving RFP alone or RFP in combination with UAS-Nup98-96 RNAi (TRiP), we confirmed effective knockdown of Nup98-96-GFP in the en-Gal4 expressing domain. Genotypes are: (left) w; Nup98-96-GFP; + (middle) w; Nup98-96-GFP/ enGal4, UAS-RFP; + (right) w; Nup98-96- GFP/ enGal4, UAS-RFP; UAS-Nup98-96 RNAi (TRiP)/+. (C) cDNA rescue constructs providing UAS-Nup98, UAS-Nup96 or UAS-Nup98-96 were tested for the ability to rescue the wing phenotypes caused by Nup98-96 RNAi. Only expression of both Nup98 and Nup96 (middle right) fully rescued posterior wing size. Note that over-expression of both Nup98 and Nup96 without RNAi also led to reduced posterior wing size (see Fig. 6). Genotypes are: (top left) w; en-Gal4, UAS-GFP/+; UAS-white RNAi (TRiP)/+ (top right) w; en-Gal4, UAS-GFP/+; UAS-Nup98-96 (middle left) w; en-Gal4, UAS-GFP/UAS-Nup98-96 RNAi (VDRC GD); UAS-white RNAi (TRiP)/+ (middle right) w; en- Gal4, UAS-GFP/UAS-Nup98-96 RNAi (VDRC GD); UASNup98-96 cDNA/+ (bottom left) w; en-Gal4, UAS-GFP/UAS-Nup98-96 RNAi (VDRC GD); UAS-Nup98 cDNA 3M/+ (bottom right) w; en-Gal4, UAS-GFP/UAS-Nup98-96 RNAi (VDRC GD); UAS-Nup96 cDNA 7M/+. (D) Two independent RNAi lines from the VDRC gave similar phenotypes to the TRiP RNAi in the larval wing. Line GD6897 is shown at wandering L3 after 72h of expression at 28°C with PH3 labeling for mitoses. Genotype: w; en-Gal4, UAS-GFP; UAS-Nup98-96 RNAi GD6897; Tub-Gal80^TS^/ +. (E) Related to EdU pulse-chase experiment in Fig. 1M. Quantification of double-labeled EdU/PH3 cells in posterior wings. Genotypes: w; en-Gal4, UAS-GFP/ + ; Tub-Gal80^TS^/ UAS-white RNAi or w; en-Gal4, UAS-GFP/ + ; Tub-Gal80^TS^/ UAS-Nup98-96 RNAi.

**Supp Fig 2:**
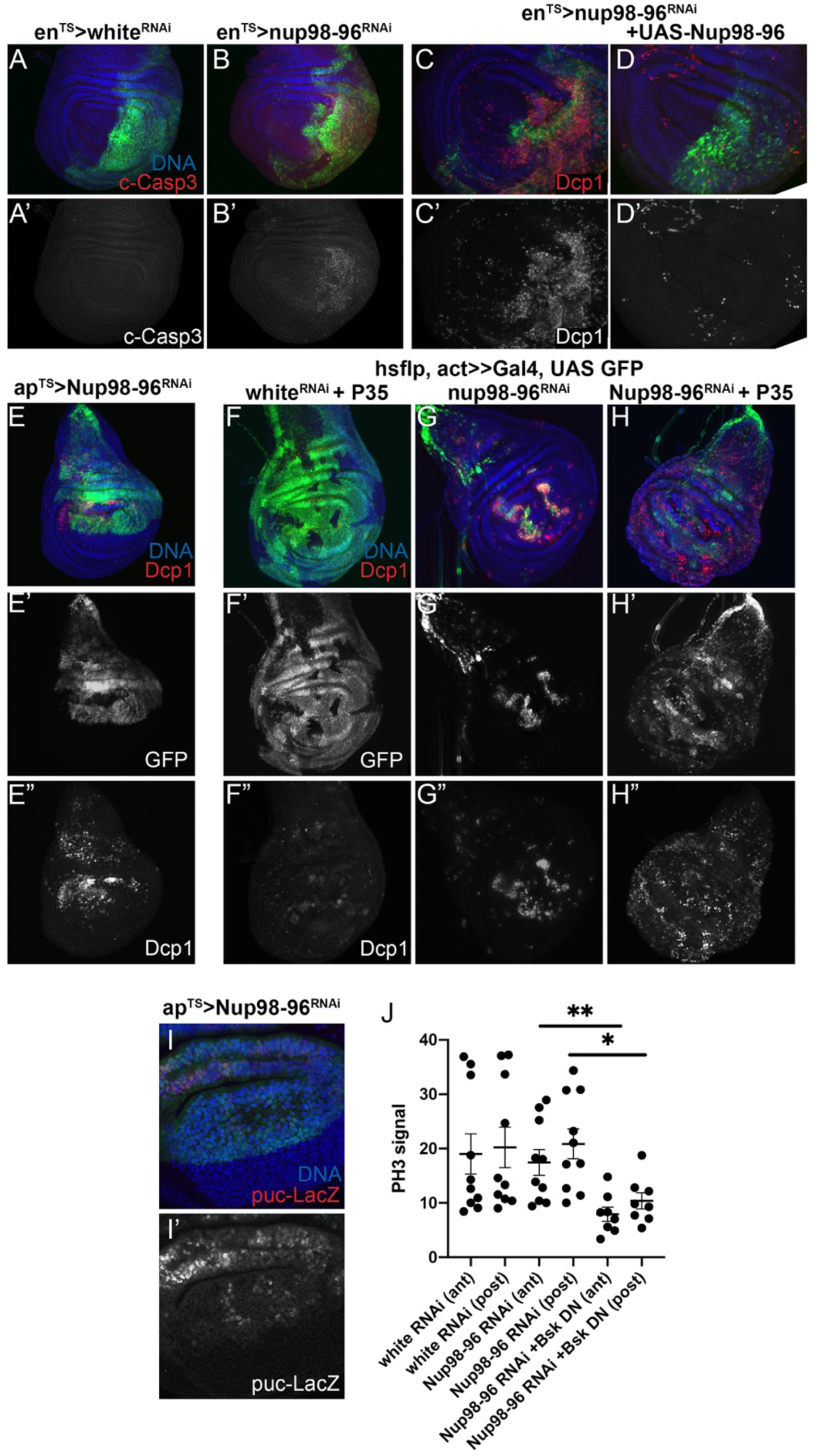
**Nup98-96 knockdown leads to apoptosis which is rescued by co-expression of UAS-Nup98-96 cDNA**. The indicated transgenes were driven by en-Gal4, with UAS GFP for 72h prior to dissection using Gal80^TS^. (A,B) Cleaved caspase 3 labeling indicates apoptosis in the posterior compartment when Nup98-96 is knocked down. (C, D) Co-expression of Nup98-96 cDNA rescues the apoptosis caused by Nup98-96 is knock-down as assessed with Dcp-1. (E) Expression of Nup98-96 RNAi in the dorsal compartment (using ap-Gal4, UAS-GFP; tub- Gal80^TS^) also leads to apoptosis. (F-H) Expression of Nup98-96 RNAi in clones throughout the wing pouch (using hs-flp with act>stop>-Gal4, UAS-GFP) also leads to apoptosis within clones, while co-expression with P35 (using hs-flp with act>stop>-Gal4, UAS-GFP, UAS-P35) suppresses apoptosis within clones and leads to apoptosis outside of clones expressing Nup98-96 RNAi. (I) Expression of Nup98-96 RNAi in the dorsal compartment (using ap-Gal4, UAS-GFP; tub-Gal80^TS^) leads to upegulation of *puc-LacZ* expression (from puc^e69^ allele), a hallmark of JNK signaling. (J) Quantifications of PH3 signal broken down by ant/post compartment. Nup98-96 RNAi and Bsk DN are expressed only in the posterior using en-Gal4, UAS-GFP; tub-Gal80^TS^. Note that overall PH3 labeling is reduced in both compartments when JNK signaling is inhibited with Bsk DN.

**Supp Fig 3:**
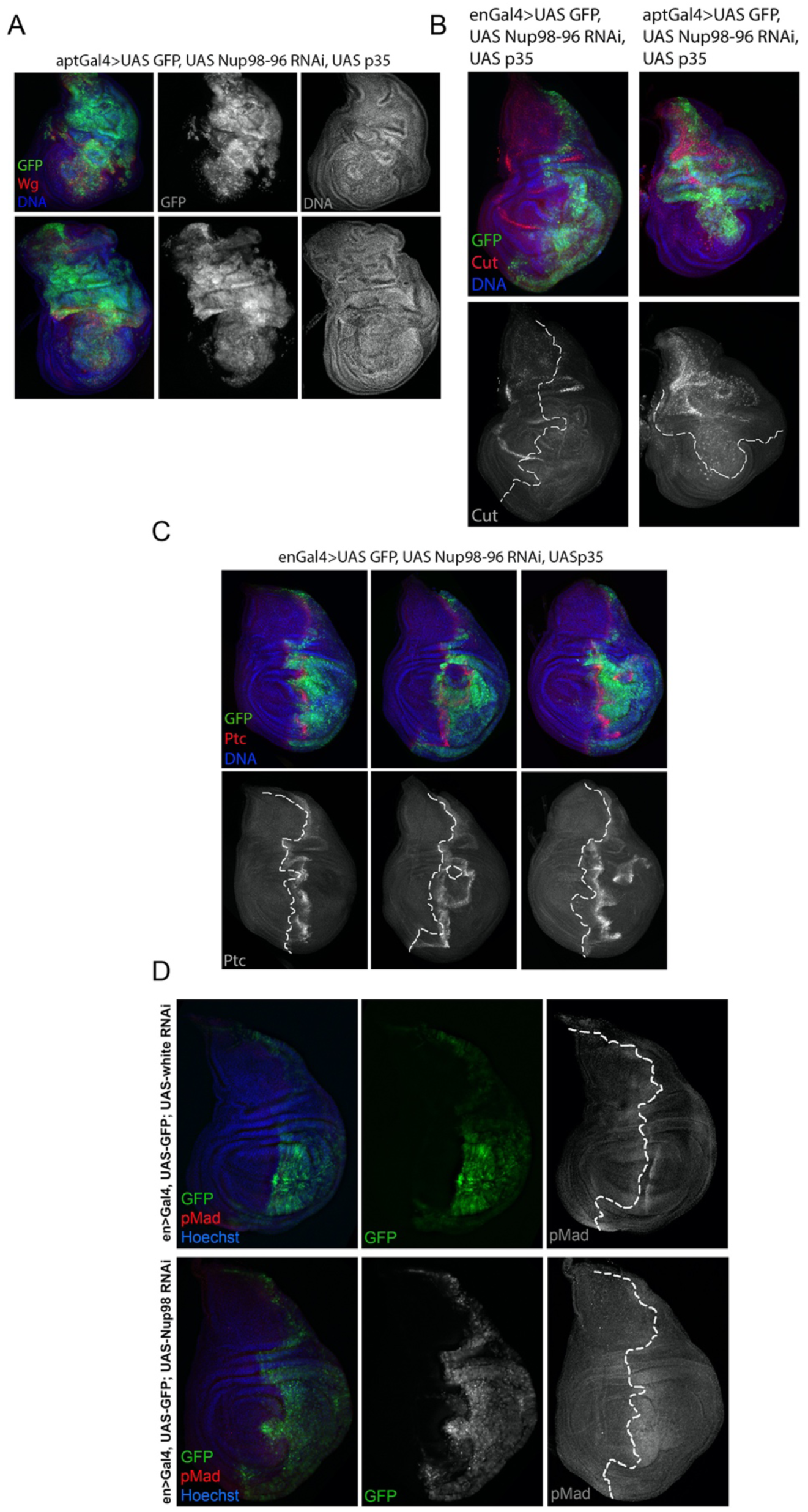
**Knockdown of Nup98-96 leads to wing overgrowth and patterning defects consistent with apoptosis-induced proliferation.** (A) Expression of Nup98-96 RNAi in the dorsal compartment with UAS-P35 for 5d (using ap- Gal4, UAS-GFP/ UAS-P35; tub-Gal80^TS^) leads to tissue folding, overgrowth and invasion across the D-V boundary as well as ectopic Wg expression. (B) Expression of Nup98-96 RNAi + P35 in the posterior compartment (with en-Gal4, left) or dorsal compartment (with ap-Gal4, right) abolishes Cut expression at the D-V boundary. (C) Expression of Nup98-96 RNAi in the posterior compartment with UAS-P35 for 4d (using en- Gal4, UAS-GFP/ UAS-P35; tub-Gal80^TS^) leads to tissue folding and invasion at the A-P boundary as well as ectopic Ptc expression demonstrating mis-patterning. (D) Normal pMad staining is shown (top) for en-Gal4, UAS-GFP; tub-Gal80^TS^ driving white RNAi. (bottom) Nup98-96 RNAi expression driven by en-Gal4, UAS-GFP; tub-Gal80^TS^ leads to broad pMad staining in the posterior compartment indicating mis-patterning.

**Supp. Fig. 4:**
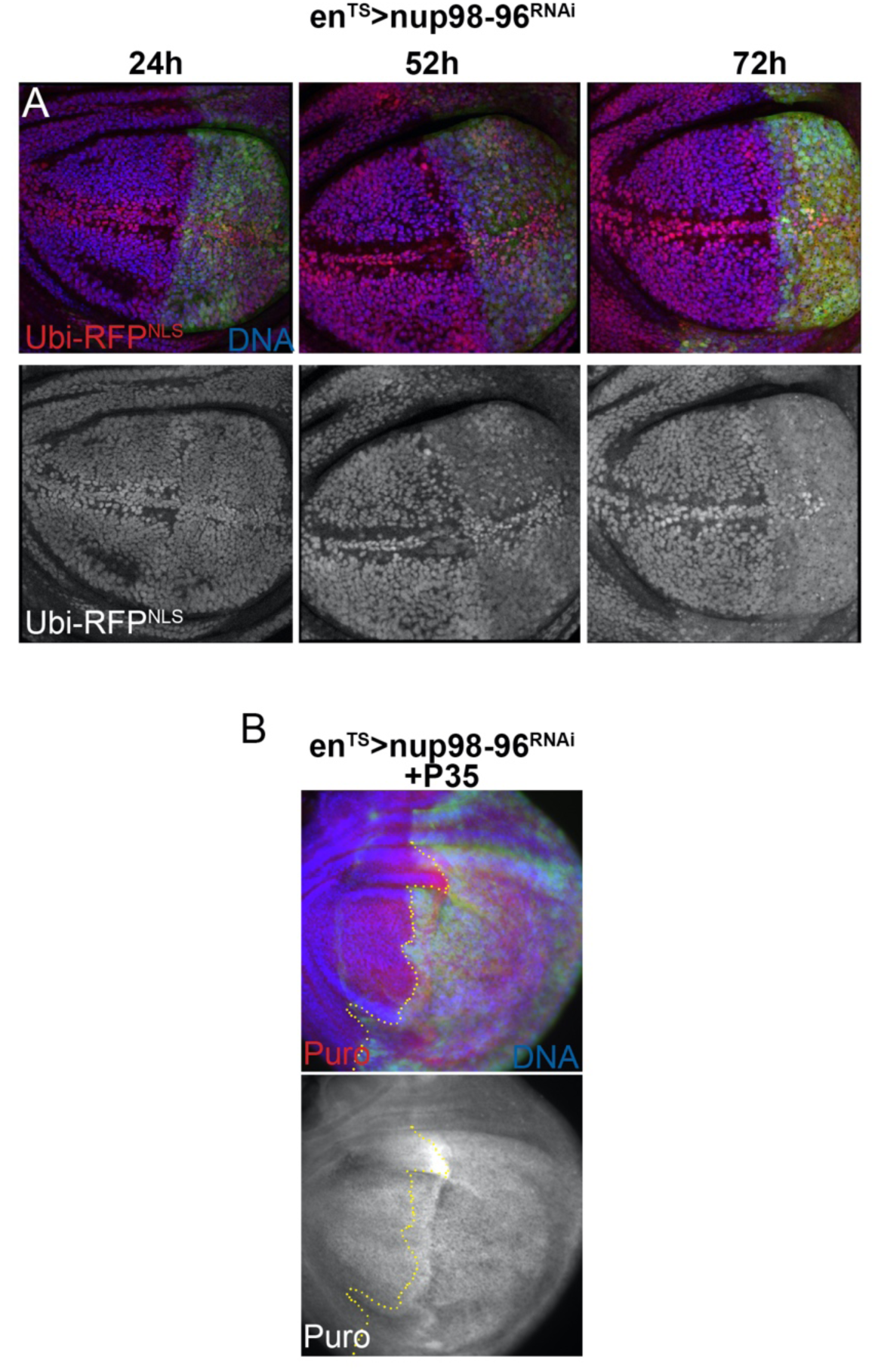
**Knockdown of Nup98-96 disrupts nucleo-cytoplasmic localization and reduces protein synthesis independent of apoptosis.** (A) Nup 98-96 RNAi was expressed in the posterior compartment using en-Gal4, UAS-GFP; tub-Gal80^TS^ in a background expressing Ubiquitin promoter driven-RFP with a nuclear localization signal (Ubi-RFP^NLS^) at 27°C to minimize cell lethality. By 52h of Nup98-96 knockdown, nuclear localization of RFP is visibly disrupted. By 72h of knockdown nuclear localization of Ubi-RFP^NLS^ is nearly abolished. (B) Co-expression of UAS-P35 with Nup 98-96 RNAi did not rescue the reduction in protein synthesis when Nup98-96 is compromised. This suggests the reduced proteins synthesis is not a consequence of apoptosis.

## Supplemental Tables

**Supplemental Table 1:** Excel file containing statistically significant gene expression changes in Nup98-96 knockdown L3 wings by RNA seq.

**Supplemental Table 2:** Excel file with the list of genes that overlap in Nup98-96 knockdown L3 wings and a wounding/regeneration program (Khan et al., 2017).

**Supplemental Table 3:** Excel file with the list of genes that overlap in Nup98-96 knockdown L3 wings and a “loser” gene expression program (Kucinski et al., 2017).

**Supplemental Table 4:** Excel file with the list of genes that overlap in Nup98-96 knockdown L3 wings and Xrp1 targets (Lee et al., 2018).

**Supplemental Table 5:** Excel file with the list of genes that overlap in Nup98-96 knockdown L3 wings and Nup98 ChIP seq assays (Pascual-Garcia et al., 2017) and Nup98 alterations with RNAseq in S2 cells (Kalverda et al., 2010).

## Notes

https://www.ncbi.nlm.nih.gov/geo/query/acc.cgi?acc=GSE152679

### Competing Interest Statement

The authors have declared no competing interest.

## References

Ahn, J. H., Davis, E. S., Daugird, T. A., Zhao, S., Quiroga, I. Y., Uryu, H., Li, J., Storey, A. J., Tsai, Y. H., Keeley, D. P., et al. (2021). Phase separation drives aberrant chromatin looping and cancer development. Nature.

Akai, N., Ohsawa, S., Sando, Y. and Igaki, T. (2021). Epithelial cell-turnover ensures robust coordination of tissue growth in Drosophila ribosomal protein mutants. PLoS Genet 17, e1009300.

Amoyel, M., Anderson, A. M. and Bach, E. A. (2014). JAK/STAT pathway dysregulation in tumors: a Drosophila perspective. Semin Cell Dev Biol 28, 96–103.

Bachi, A., Braun, I. C., Rodrigues, J. P., Pante, N., Ribbeck, K., von Kobbe, C., Kutay, U., Wilm, M., Gorlich, D., Carmo-Fonseca, M., et al. (2000). The C-terminal domain of TAP interacts with the nuclear pore complex and promotes export of specific CTE-bearing RNA substrates. RNA 6, 136–158.

Bandura, J. L., Jiang, H., Nickerson, D. W. and Edgar, B. A. (2013). The molecular chaperone Hsp90 is required for cell cycle exit in Drosophila melanogaster. PLoS Genet 9, e1003835.

Baonza, A., Roch, F. and Martin-Blanco, E. (2000). DER signaling restricts the boundaries of the wing field during Drosophila development. Proc Natl Acad Sci U S A 97, 7331–7335.

Baril, C., Gavory, G., Bidla, G., Knaevelsrud, H., Sauvageau, G. and Therrien, M. (2017). Human NUP98-HOXA9 promotes hyperplastic growth of hematopoietic tissues in Drosophila. Dev Biol 421, 16–26.

Baumgartner, M. E., Dinan, M. P., Langton, P. F., Kucinski, I. and Piddini, E. (2021). Proteotoxic stress is a driver of the loser status and cell competition. Nat Cell Biol 23, 136–146.

Bergantinos, C., Corominas, M. and Serras, F. (2010). Cell death-induced regeneration in wing imaginal discs requires JNK signalling. Development 137, 1169–1179.

Braune, E. B. and Lendahl, U. (2016). Notch -- a goldilocks signaling pathway in disease and cancer therapy. Discov Med 21, 189–196.

Brodsky, M. H., Nordstrom, W., Tsang, G., Kwan, E., Rubin, G. M. and Abrams, J. M. (2000). Drosophila p53 binds a damage response element at the reaper locus. Cell 101, 103–113.

Brown, J. B., Boley, N., Eisman, R., May, G. E., Stoiber, M. H., Duff, M. O., Booth, B. W., Wen, J., Park, S., Suzuki, A. M., et al. (2014). Diversity and dynamics of the Drosophila transcriptome. Nature 512, 393–399.

Buttitta, L. A., Katzaroff, A. J. and Edgar, B. A. (2010). A robust cell cycle control mechanism limits E2F-induced proliferation of terminally differentiated cells in vivo. J Cell Biol 189, 981–996.

Buttitta, L. A., Katzaroff, A. J., Perez, C. L., de la Cruz, A. and Edgar, B. A. (2007). A double-assurance mechanism controls cell cycle exit upon terminal differentiation in Drosophila. Dev Cell 12, 631–643.

Capelson, M., Liang, Y., Schulte, R., Mair, W., Wagner, U. and Hetzer, M. W. (2010). Chromatin-bound nuclear pore components regulate gene expression in higher eukaryotes. Cell 140, 372–383.

Chaichanit, N., Wonglapsuwan, M. and Chotigeat, W. (2018). Ribosomal protein L10A and signaling pathway. Gene 674, 170–177.

Chakraborty, P., Wang, Y., Wei, J. H., van Deursen, J., Yu, H., Malureanu, L., Dasso, M., Forbes, D. J., Levy, D. E., Seemann, J., et al. (2008). Nucleoporin levels regulate cell cycle progression and phase-specific gene expression. Dev Cell 15, 657–667.

Cheng, G., Liu, F., Asai, T., Lai, F., Man, N., Xu, H., Chen, S., Greenblatt, S., Hamard, P. J., Ando, K., et al. (2017). Loss of p300 accelerates MDS-associated leukemogenesis. Leukemia 31, 1382–1390.

Choi, C. W., Chung, Y. J., Slape, C. and Aplan, P. D. (2008). Impaired differentiation and apoptosis of hematopoietic precursors in a mouse model of myelodysplastic syndrome. Haematologica 93, 1394–1397.

Colombani, J., Andersen, D. S. and Leopold, P. (2012). Secreted peptide Dilp8 coordinates Drosophila tissue growth with developmental timing. Science 336, 582–585.

de Celis, J. F., Garcia-Bellido, A. and Bray, S. J. (1996). Activation and function of Notch at the dorsal-ventral boundary of the wing imaginal disc. Development 122, 359–369.

Deliu, L. P., Ghosh, A. and Grewal, S. S. (2017). Investigation of protein synthesis in Drosophila larvae using puromycin labelling. Biol Open 6, 1229–1234.

Dimova, D. K., Stevaux, O., Frolov, M. V. and Dyson, N. J. (2003). Cell cycle-dependent and cell cycle-independent control of transcription by the Drosophila E2F/RB pathway. Genes Dev 17, 2308–2320.

Duman-Scheel, M., Johnston, L. A. and Du, W. (2004). Repression of dMyc expression by Wingless promotes Rbf-induced G1 arrest in the presumptive Drosophila wing margin. Proc Natl Acad Sci U S A 101, 3857–3862.

Dvorak, H. F. (1986). Tumors: wounds that do not heal. Similarities between tumor stroma generation and wound healing. N Engl J Med 315, 1650–1659.

Fahrenkrog, B., Martinelli, V., Nilles, N., Fruhmann, G., Chatel, G., Juge, S., Sauder, U., Di Giacomo, D., Mecucci, C. and Schwaller, J. (2016). Expression of Leukemia-Associated Nup98 Fusion Proteins Generates an Aberrant Nuclear Envelope Phenotype. PLoS One 11, e0152321.

Faria, A. M., Levay, A., Wang, Y., Kamphorst, A. O., Rosa, M. L., Nussenzveig, D. R., Balkan, W., Chook, Y. M., Levy, D. E. and Fontoura, B. M. (2006). The nucleoporin Nup96 is required for proper expression of interferon-regulated proteins and functions. Immunity 24, 295–304.

Flegel, K., Grushko, O., Bolin, K., Griggs, E. and Buttitta, L. (2016). Roles for the Histone Modifying and Exchange Complex NuA4 in Cell Cycle Progression in Drosophila melanogaster. Genetics 203, 1265–1281.

Fogarty, C. E. and Bergmann, A. (2017). Killers creating new life: caspases drive apoptosis-induced proliferation in tissue repair and disease. Cell Death Differ 24, 1390–1400.

Fontoura, B. M., Blobel, G. and Matunis, M. J. (1999). A conserved biogenesis pathway for nucleoporins: proteolytic processing of a 186-kilodalton precursor generates Nup98 and the novel nucleoporin, Nup96. J Cell Biol 144, 1097–1112.

Forrester, A. M., Grabher, C., McBride, E. R., Boyd, E. R., Vigerstad, M. H., Edgar, A., Kai, F. B., Da’as, S. I., Payne, E., Look, A. T., et al. (2011). NUP98-HOXA9-transgenic zebrafish develop a myeloproliferative neoplasm and provide new insight into mechanisms of myeloid leukaemogenesis. Br J Haematol 155, 167–181.

Franks, T. M. and Hetzer, M. W. (2013). The role of Nup98 in transcription regulation in healthy and diseased cells. Trends Cell Biol 23, 112–117.

Gabriele, L., Phung, J., Fukumoto, J., Segal, D., Wang, I. M., Giannakakou, P., Giese, N. A., Ozato, K. and Morse, H. C., 3rd (1999). Regulation of apoptosis in myeloid cells by interferon consensus sequence-binding protein. J Exp Med 190, 411–421.

Garelli, A., Gontijo, A. M., Miguela, V., Caparros, E. and Dominguez, M. (2012). Imaginal discs secrete insulin-like peptide 8 to mediate plasticity of growth and maturation. Science 336, 579–582.

Gleizes, P. E., Noaillac-Depeyre, J., Leger-Silvestre, I., Teulieres, F., Dauxois, J. Y., Pommet, D., Azum-Gelade, M. C. and Gas, N. (2001). Ultrastructural localization of rRNA shows defective nuclear export of preribosomes in mutants of the Nup82p complex. J Cell Biol 155, 923–936.

Gough, S. M., Slape, C. I. and Aplan, P. D. (2011). NUP98 gene fusions and hematopoietic malignancies: common themes and new biologic insights. Blood 118, 6247–6257.

Grewal, S. S. (2009). Insulin/TOR signaling in growth and homeostasis: a view from the fly world. Int J Biochem Cell Biol 41, 1006–1010.

Griffis, E. R., Altan, N., Lippincott-Schwartz, J. and Powers, M. A. (2002). Nup98 is a mobile nucleoporin with transcription-dependent dynamics. Mol Biol Cell 13, 1282–1297.

Gurevich, R. M., Rosten, P. M., Schwieger, M., Stocking, C. and Humphries, R. K. (2006). Retroviral integration site analysis identifies ICSBP as a collaborating tumor suppressor gene in NUP98-TOP1-induced leukemia. Exp Hematol 34, 1192–1201.

Halder, G., Polaczyk, P., Kraus, M. E., Hudson, A., Kim, J., Laughon, A. and Carroll, S. (1998). The Vestigial and Scalloped proteins act together to directly regulate wing-specific gene expression in Drosophila. Genes Dev 12, 3900–3909.

Hay, B. A., Wolff, T. and Rubin, G. M. (1994). Expression of baculovirus P35 prevents cell death in Drosophila. Development 120, 2121–2129.

Herranz, H., Perez, L., Martin, F. A. and Milan, M. (2008). A Wingless and Notch double-repression mechanism regulates G1-S transition in the Drosophila wing. EMBO J 27, 1633–1645.

Herrera, S. C., Martin, R. and Morata, G. (2013). Tissue homeostasis in the wing disc of Drosophila melanogaster: immediate response to massive damage during development. PLoS Genet 9, e1003446.

Hu, L., Huang, W., Hjort, E. E., Bei, L., Platanias, L. C. and Eklund, E. A. (2016). The Interferon Consensus Sequence Binding Protein (Icsbp/Irf8) Is Required for Termination of Emergency Granulopoiesis. J Biol Chem 291, 4107–4120.

Jeganathan, K. B., Baker, D. J. and van Deursen, J. M. (2006). Securin associates with APCCdh1 in prometaphase but its destruction is delayed by Rae1 and Nup98 until the metaphase/anaphase transition. Cell Cycle 5, 366–370.

Jeganathan, K. B., Malureanu, L. and van Deursen, J. M. (2005). The Rae1-Nup98 complex prevents aneuploidy by inhibiting securin degradation. Nature 438, 1036–1039.

Ji, Z., Kiparaki, M., Folgado, V., Kumar, A., Blanco, J., Rimesso, G., Chuen, J., Liu, Y., Zheng, D. and Baker, N. E. (2019). Drosophila RpS12 controls translation, growth, and cell competition through Xrp1. PLoS Genet 15, e1008513.

Johnson, A. W., Lund, E. and Dahlberg, J. (2002). Nuclear export of ribosomal subunits. Trends Biochem Sci 27, 580–585.

Johnston, L. A. and Edgar, B. A. (1998). Wingless and Notch regulate cell-cycle arrest in the developing Drosophila wing. Nature 394, 82–84.

Joyce, J. A. and Schofield, P. N. (1998). Genomic imprinting and cancer. Mol Pathol 51, 185–190.

Kalverda, B., Pickersgill, H., Shloma, V. V. and Fornerod, M. (2010). Nucleoporins directly stimulate expression of developmental and cell-cycle genes inside the nucleoplasm. Cell 140, 360–371.

Karin, M. and Clevers, H. (2016). Reparative inflammation takes charge of tissue regeneration. Nature 529, 307–315.

Katsuyama, T., Comoglio, F., Seimiya, M., Cabuy, E. and Paro, R. (2015). During Drosophila disc regeneration, JAK/STAT coordinates cell proliferation with Dilp8-mediated developmental delay. Proc Natl Acad Sci U S A 112, E2327–2336.

Khan, S. J., Abidi, S. N. F., Skinner, A., Tian, Y. and Smith-Bolton, R. K. (2017). The Drosophila Duox maturation factor is a key component of a positive feedback loop that sustains regeneration signaling. PLoS Genet 13, e1006937.

Kim, J., Sebring, A., Esch, J. J., Kraus, M. E., Vorwerk, K., Magee, J. and Carroll, S. B. (1996). Integration of positional signals and regulation of wing formation and identity by Drosophila vestigial gene. Nature 382, 133–138.

Kucinski, I., Dinan, M., Kolahgar, G. and Piddini, E. (2017). Chronic activation of JNK JAK/STAT and oxidative stress signalling causes the loser cell status. Nat Commun 8, 136.

Kulshammer, E., Mundorf, J., Kilinc, M., Frommolt, P., Wagle, P. and Uhlirova, M. (2015). Interplay among Drosophila transcription factors Ets21c, Fos and Ftz-F1 drives JNK-mediated tumor malignancy. Dis Model Mech 8, 1279–1293.

Lam, D. H. and Aplan, P. D. (2001). NUP98 gene fusions in hematologic malignancies. Leukemia 15, 1689–1695.

Lapik, Y. R., Fernandes, C. J., Lau, L. F. and Pestov, D. G. (2004). Physical and functional interaction between Pes1 and Bop1 in mammalian ribosome biogenesis. Mol Cell 15, 17–29.

Law, C. W., Chen, Y., Shi, W. and Smyth, G. K. (2014). voom: Precision weights unlock linear model analysis tools for RNA-seq read counts. Genome Biol 15, R29.

Lee, C. H., Kiparaki, M., Blanco, J., Folgado, V., Ji, Z., Kumar, A., Rimesso, G. and Baker, N. E. (2018). A Regulatory Response to Ribosomal Protein Mutations Controls Translation, Growth, and Cell Competition. Dev Cell 46, 807.

Liao, Y., Smyth, G. K. and Shi, W. (2014). featureCounts: an efficient general purpose program for assigning sequence reads to genomic features. Bioinformatics 30, 923–930.

Lin, Y. W., Slape, C., Zhang, Z. and Aplan, P. D. (2005). NUP98-HOXD13 transgenic mice develop a highly penetrant, severe myelodysplastic syndrome that progresses to acute leukemia. Blood 106, 287–295.

Lo, K. Y., Li, Z., Bussiere, C., Bresson, S., Marcotte, E. M. and Johnson, A. W. (2010). Defining the pathway of cytoplasmic maturation of the 60S ribosomal subunit. Mol Cell 39, 196–208.

Lockhead, S., Moskaleva, A., Kamenz, J., Chen, Y., Kang, M., Reddy, A. R., Santos, S. D. M. and Ferrell, J. E., Jr. (2020). The Apparent Requirement for Protein Synthesis during G2 Phase Is due to Checkpoint Activation. Cell Rep 32, 107901.

Ma, C., Wu, S., Li, N., Chen, Y., Yan, K., Li, Z., Zheng, L., Lei, J., Woolford, J. L., Jr. and Gao, N. (2017). Structural snapshot of cytoplasmic pre-60S ribosomal particles bound by Nmd3, Lsg1, Tif6 and Reh1. Nat Struct Mol Biol 24, 214–220.

Marygold, S. J., Roote, J., Reuter, G., Lambertsson, A., Ashburner, M., Millburn, G. H., Harrison, P. M., Yu, Z., Kenmochi, N., Kaufman, T. C., et al. (2007). The ribosomal protein genes and Minute loci of Drosophila melanogaster. Genome Biol 8, R216.

McEwen, D. G. and Peifer, M. (2005). Puckered, a Drosophila MAPK phosphatase, ensures cell viability by antagonizing JNK-induced apoptosis. Development 132, 3935–3946.

Mendes, A., Juhlen, R., Bousbata, S. and Fahrenkrog, B. (2020). Disclosing the Interactome of Leukemogenic NUP98-HOXA9 and SET-NUP214 Fusion Proteins Using a Proteomic Approach. Cells 9.

Mesquita, D., Dekanty, A. and Milan, M. (2010). A dp53-dependent mechanism involved in coordinating tissue growth in Drosophila. PLoS Biol 8, e1000566.

Mondal, B. C., Shim, J., Evans, C. J. and Banerjee, U. (2014). Pvr expression regulators in equilibrium signal control and maintenance of Drosophila blood progenitors. Elife 3, e03626.

Moroianu, J., Hijikata, M., Blobel, G. and Radu, A. (1995). Mammalian karyopherin alpha 1 beta and alpha 2 beta heterodimers: alpha 1 or alpha 2 subunit binds nuclear localization signal and beta subunit interacts with peptide repeat-containing nucleoporins. Proc Natl Acad Sci U S A 92, 6532–6536.

Moy, T. I. and Silver, P. A. (2002). Requirements for the nuclear export of the small ribosomal subunit. J Cell Sci 115, 2985–2995.

Musalgaonkar, S., Black, J. J. and Johnson, A. W. (2019). The L1 stalk is required for efficient export of nascent large ribosomal subunits in yeast. RNA 25, 1549–1560.

Muzzopappa, M., Murcia, L. and Milan, M. (2017). Feedback amplification loop drives malignant growth in epithelial tissues. Proc Natl Acad Sci U S A 114, E7291–E7300.

Narbonne-Reveau, K. and Maurange, C. (2019). Developmental regulation of regenerative potential in Drosophila by ecdysone through a bistable loop of ZBTB transcription factors. PLoS Biol 17, e3000149.

Neufeld, T. P., de la Cruz, A. F., Johnston, L. A. and Edgar, B. A. (1998). Coordination of growth and cell division in the Drosophila wing. Cell 93, 1183–1193.

Neumann, C. J. and Cohen, S. M. (1997). Long-range action of Wingless organizes the dorsal-ventral axis of the Drosophila wing. Development 124, 871–880.

O’Keefe, D. D., Thomas, S. R., Bolin, K., Griggs, E., Edgar, B. A. and Buttitta, L. A. (2012). Combinatorial control of temporal gene expression in the Drosophila wing by enhancers and core promoters. BMC Genomics 13, 498.

Oeffinger, M., Dlakic, M. and Tollervey, D. (2004). A pre-ribosome-associated HEAT-repeat protein is required for export of both ribosomal subunits. Genes Dev 18, 196–209.

Pascual-Garcia, P., Debo, B., Aleman, J. R., Talamas, J. A., Lan, Y., Nguyen, N. H., Won, K. J. and Capelson, M. (2017). Metazoan Nuclear Pores Provide a Scaffold for Poised Genes and Mediate Induced Enhancer-Promoter Contacts. Mol Cell 66, 63–76 e66.

Pascual-Garcia, P., Jeong, J. and Capelson, M. (2014). Nucleoporin Nup98 associates with Trx/MLL and NSL histone-modifying complexes and regulates Hox gene expression. Cell Rep 9, 433–442.

Perez-Garijo, A., Shlevkov, E. and Morata, G. (2009). The role of Dpp and Wg in compensatory proliferation and in the formation of hyperplastic overgrowths caused by apoptotic cells in the Drosophila wing disc. Development 136, 1169–1177.

Pinal, N., Martin, M., Medina, I. and Morata, G. (2018). Short-term activation of the Jun N-terminal kinase pathway in apoptosis-deficient cells of Drosophila induces tumorigenesis. Nat Commun 9, 1541.

Reber, A., Lehner, C. F. and Jacobs, H. W. (2006). Terminal mitoses require negative regulation of Fzr/Cdh1 by Cyclin A, preventing premature degradation of mitotic cyclins and String/Cdc25. Development 133, 3201–3211.

Romero-Pozuelo, J., Demetriades, C., Schroeder, P. and Teleman, A. A. (2017). CycD/Cdk4 and Discontinuities in Dpp Signaling Activate TORC1 in the Drosophila Wing Disc. Dev Cell 42, 376–387 e375.

Romero-Pozuelo, J., Figlia, G., Kaya, O., Martin-Villalba, A. and Teleman, A. A. (2020). Cdk4 and Cdk6 Couple the Cell-Cycle Machinery to Cell Growth via mTORC1. Cell Rep 31, 107504.

Rosenblum, J. S. and Blobel, G. (1999). Autoproteolysis in nucleoporin biogenesis. Proc Natl Acad Sci U S A 96, 11370–11375.

Ruggiero, R., Kale, A., Thomas, B. and Baker, N. E. (2012). Mitosis in neurons: Roughex and APC/C maintain cell cycle exit to prevent cytokinetic and axonal defects in Drosophila photoreceptor neurons. PLoS Genet 8, e1003049.

Sabri, N., Roth, P., Xylourgidis, N., Sadeghifar, F., Adler, J. and Samakovlis, C. (2007). Distinct functions of the Drosophila Nup153 and Nup214 FG domains in nuclear protein transport. J Cell Biol 178, 557–565.

Sarov, M., Barz, C., Jambor, H., Hein, M. Y., Schmied, C., Suchold, D., Stender, B., Janosch, S., K, J. V., Krishnan, R. T., et al. (2016). A genome-wide resource for the analysis of protein localisation in Drosophila. Elife 5, e12068.

Schmidt, H. B. and Gorlich, D. (2015). Nup98 FG domains from diverse species spontaneously phase-separate into particles with nuclear pore-like permselectivity. Elife 4.

Schuster, K. J. and Smith-Bolton, R. K. (2015). Taranis Protects Regenerating Tissue from Fate Changes Induced by the Wound Response in Drosophila. Dev Cell 34, 119–128.

Shi, Z., Fujii, K., Kovary, K. M., Genuth, N. R., Rost, H. L., Teruel, M. N. and Barna, M. (2017). Heterogeneous Ribosomes Preferentially Translate Distinct Subpools of mRNAs Genome-wide. Mol Cell 67, 71–83 e77.

Simon, D. N. and Rout, M. P. (2014). Cancer and the nuclear pore complex. Adv Exp Med Biol 773, 285–307.

Singer, S., Zhao, R., Barsotti, A. M., Ouwehand, A., Fazollahi, M., Coutavas, E., Breuhahn, K., Neumann, O., Longerich, T., Pusterla, T., et al. (2012). Nuclear pore component Nup98 is a potential tumor suppressor and regulates posttranscriptional expression of select p53 target genes. Mol Cell 48, 799–810.

Slape, C., Liu, L. Y., Beachy, S. and Aplan, P. D. (2008). Leukemic transformation in mice expressing a NUP98-HOXD13 transgene is accompanied by spontaneous mutations in Nras, Kras, and Cbl. Blood 112, 2017–2019.

Smith-Bolton, R. K., Worley, M. I., Kanda, H. and Hariharan, I. K. (2009). Regenerative growth in Drosophila imaginal discs is regulated by Wingless and Myc. Dev Cell 16, 797–809.

Sullivan, W. A., M; Hawley, RS (2000). *Drosophila Protocols*: Cold Spring Harbor Press.

Sun, D. and Buttitta, L. (2015). Protein phosphatase 2A promotes the transition to G0 during terminal differentiation in Drosophila. Development 142, 3033–3045.

Tanaka-Matakatsu, M., Thomas, B. J. and Du, W. (2007). Mutation of the Apc1 homologue shattered disrupts normal eye development by disrupting G1 cell cycle arrest and progression through mitosis. Dev Biol 309, 222–235.

Thomas, A., Lee, P. J., Dalton, J. E., Nomie, K. J., Stoica, L., Costa-Mattioli, M., Chang, P., Nuzhdin, S., Arbeitman, M. N. and Dierick, H. A. (2012). A versatile method for cell-specific profiling of translated mRNAs in Drosophila. PLoS One 7, e40276.

Uhlirova, M. and Bohmann, D. (2006). JNK- and Fos-regulated Mmp1 expression cooperates with Ras to induce invasive tumors in Drosophila. EMBO J 25, 5294–5304.

Verghese, S. and Su, T. T. (2017). STAT, Wingless, and Nurf-38 determine the accuracy of regeneration after radiation damage in Drosophila. PLoS Genet 13, e1007055.

Walther, T. C., Alves, A., Pickersgill, H., Loiodice, I., Hetzer, M., Galy, V., Hulsmann, B. B., Kocher, T., Wilm, M., Allen, T., et al. (2003). The conserved Nup107-160 complex is critical for nuclear pore complex assembly. Cell 113, 195–206.

Wild, T., Horvath, P., Wyler, E., Widmann, B., Badertscher, L., Zemp, I., Kozak, K., Csucs, G., Lund, E. and Kutay, U. (2010). A protein inventory of human ribosome biogenesis reveals an essential function of exportin 5 in 60S subunit export. PLoS Biol 8, e1000522.

Williams, J. A., Bell, J. B. and Carroll, S. B. (1991). Control of Drosophila wing and haltere development by the nuclear vestigial gene product. Genes Dev 5, 2481–2495.

Williams, J. A., Paddock, S. W. and Carroll, S. B. (1993). Pattern formation in a secondary field: a hierarchy of regulatory genes subdivides the developing Drosophila wing disc into discrete subregions. Development 117, 571–584.

Wonglapsuwan, M., Chotigeat, W., Timmons, A. and McCall, K. (2011). RpL10A regulates oogenesis progression in the banana prawn Fenneropenaeus merguiensis and Drosophila melanogaster. Gen Comp Endocrinol 173, 356–363.

Worley, M. I., Alexander, L. A. and Hariharan, I. K. (2018). CtBP impedes JNK- and Upd/STAT-driven cell fate misspecifications in regenerating Drosophila imaginal discs. Elife 7.

Wu, X., Kasper, L. H., Mantcheva, R. T., Mantchev, G. T., Springett, M. J. and van Deursen, J. M. (2001). Disruption of the FG nucleoporin NUP98 causes selective changes in nuclear pore complex stoichiometry and function. Proc Natl Acad Sci U S A 98, 3191–3196.

Xu, S. and Powers, M. A. (2009). Nuclear pore proteins and cancer. Semin Cell Dev Biol 20, 620–630.

Zecca, M. and Struhl, G. (2010). A feed-forward circuit linking wingless, fat-dachsous signaling, and the warts-hippo pathway to Drosophila wing growth. PLoS Biol 8, e1000386.

